# Nucleus accumbens D2-expressing neurons: Balancing reward and licking disruption through rhythmic optogenetic stimulation

**DOI:** 10.1101/2024.11.04.621937

**Authors:** Nikte Requejo-Mendoza, José-Antonio Arias-Montaño, Ranier Gutierrez

**Author notes:** Corresponding author (RG).

## Abstract

Dopamine D1 receptor-expressing neurons in the nucleus accumbens (NAc) are known to be critical for processing reward and regulating food intake. However, the role of D2-expressing neurons in this region remains less understood. This study employed optogenetic manipulations to investigate the role of NAc D2-expressing neurons in reward processing and sucrose consumption. Optogenetic activation of these neurons decreased sucrose preference (at 20 Hz), disrupted licking patterns (particularly at 8 and 20 Hz), and increased self-stimulation. Conversely, synchronizing stimulation with the animal’s licking rhythm mitigated licking disruption and even increased sucrose intake, suggesting a rewarding effect. Furthermore, 20 Hz stimulation (but not 8 Hz) induced place preference in a real-time place preference (RTPP) test. In contrast, inhibiting D2 neurons produced a negative hedonic state, although not reaching a complete aversion, influencing food choices in specific contexts. For instance, while the RTPP test per se was not sensitive enough to observe place aversion when mice could choose between consuming a high-fat diet (HFD) pellet in a context associated with or without inhibition of D2 neurons, they preferred to consume HFD on the non-inhibited side. This suggests that the palatability of HFD can unmask (but also overshadow) the negative hedonic state associated with D2 neuron inhibition. A negative reinforcement paradigm further confirmed the active avoidance behavior induced by D2 neuron inhibition. In conclusion, NAc D2 neuron inhibition induces a negative hedonic state, while activation has a dual effect—it is rewarding yet disrupts licking behavior—highlighting its complex role in reward and consummatory behavior. Importantly, self-paced stimulation, where the animal controls the timing of the stimulation by its licking behavior, offers a more efficient and natural approach for stimulating NAc activity.

## Introduction

The nucleus accumbens (NAc) is known to be involved in reward processing and food intake.[1–3] While the role of dopamine D1 receptor-expressing neurons in the NAc has been extensively studied, the role of D2 neurons remains unclear.[4,5] D1 medium spiny neurons (MSNs) in the NAc are well-established as key players in processing rewards and regulating food intake.[6,7] For instance, D1 neuron inhibition permits food consumption, whereas activation suppresses licking behavior, a fundamental component of the feeding-gating hypothesis of the NAc, particularly of the shell region.[8,9] In contrast, feeding has been reported to have no modulatory effect on D2 receptor-expressing neurons.[9,10] Accordingly, a recent study using rabies virus has traced that D1 NAc neurons receive direct inputs from taste bud cells on the tongue and send direct projections to the reticular formation in which the Central Pattern Generator (CPG) for rhythmic licking is controlled. In contrast, the same study showed that D2 neurons are not directly connected with the CPG controlling tongue movements.[11]

Beyond its involvement in food consumption, the NAc has been recognized as a crucial player in reward processing since the groundbreaking work of James Olds (1956) on the brain pleasure centers.[12] This role is also supported by evidence that mice will self-stimulate to activate all glutamatergic inputs to the NAc shell, [13,14] indicating the capability of glutamate to modulate reward and even suppress feeding behavior, presumably by influencing the activity of either D1 or D2 receptor-expressing neurons.[14] The classic view of D1- and D2-expressing neurons posited opposing roles in feeding[15] and in reward, D1 neuron activation being rewarding and D2 activation aversive, with inhibition leading to the opposite effects.[16] However, current research suggests a more complex dynamic. While D1-expressing neurons predominantly drive positive reinforcement and reward behaviors,[16–18] D2-expressing neurons appear to have more complex, bidirectional effects, depending on the context and neural stimulation patterns.[4] Soares-Cunha work further challenged the classic view, demonstrating that brief (1 s, 40 Hz) optogenetic stimulation of D1- or D2-expressing neurons is rewarding, whereas prolonged stimulation (60 s) elicits aversion.[5] This highlights that the rewarding or aversive nature of D1 and D2 neuron activity hinges on stimulation parameters.[5] Thus, while the role of NAc D1-expressing neurons in reward is more established,[19,20] the contribution of D2-expressing neurons yields variable results across different behaviors and experimental setups.[4,5,11,17]

This study employed optogenetic manipulations to investigate the specific role of NAc D2-expressing neurons in reward processing and consummatory behavior related to sucrose and HFD intake. By exploring multiple frequency-dependent parameters for optogenetic manipulation, we aimed to provide further insights into how D2 neurons fine-tune reward processing and consummatory behaviors. Our data demonstrate the complex nature of NAc D2 neuron activation, revealing its dual role in promoting reward while disrupting certain aspects of consummatory behavior. In contrast, NAc D2 neuron inhibition induces a negative hedonic value. Furthermore, our findings highlight the importance of stimulation parameters in understanding NAc D2 neuron function

## Materials and methods

### Ethical Considerations

Post-surgical animal care included administering analgesics (lidocaine hydrochloride, 5 mg/kg) and antibiotics (enrofloxacin, 5 mg/kg). Animals were monitored daily for signs of pain or distress, including weight loss, immobility, or signs of infection. No animals required early euthanasia during the study. At the end of the experiments, animals were humanely euthanized via an overdose of sodium pentobarbital, followed by transcardial perfusion with phosphate-buffered saline and 4% paraformaldehyde. All experimental procedures were approved by the Cinvestav Animal Care and Use Committee (CINVESTAV) under permit number [0080–14] and conducted the Guide for the Care and Use of Laboratory Animals of the National Institutes of Health.

### Animals

Drd2-Cre (B6.FVB(Cg)-Tg(Drd2-cre)ER44Gsat/Mmcd) mice of both sexes (7-10 weeks old, 20-25 g at the beginning of the experiment; MMRC, stock number: 032108) were single-housed in a temperature-controlled room (22 ± 1°C) with a 12-h light/dark cycle. Mice had *ad libitum* access to a standard chow diet (PicoLab Rodent Diet 20, MO, USA) and water unless otherwise specified for a particular behavioral paradigm. After surgery, mice were single-housed and maintained in a temperature-controlled environment with a 12-h light/dark cycle.

### Viral constructs

For cell-type-specific manipulation of D2-expressing neurons in the NAc, mice were injected with Cre-dependent adeno-associated viral (AAV) vectors carrying genes encoding Channelrhodopsin (ChR2) for optogenetic activation and Archaerhodopsin (ArchT) for neuronal inhibition, as well as control vectors like tdTomato. AAV vectors were obtained from Addgene (www.addgene.org): ChR2-eYFP (AAV5-EfIa-DIOhChR2(E123T/T159C)-EYFP, #35509) at a titer of 1 x 1013 vector genome/ml (vg/ml); ArchT-tdTomato (AAV5/FLEX-ArchT-tdTomato, #28305) at 7.0 x 1012 vg/ml; eYFP-vector (AAV5-EfIa-DIO EYFP, #27056) at 1.0 x 1013 vg/ml; and AAV5 pCAG-FLEX-tdTomato-WPRE (#51503) at 3.8 x 1013 vg/ml. Viruses were aliquoted and stored at −80°C until use.

### Stereotaxic surgery, viral infection, and fiber implantation

Mice were anesthetized with 2% isoflurane (flow rate 0.5 L/min), and before surgery, mice received a subcutaneous analgesic (lidocaine hydrochloride; 5 mg/kg) on the scalp and topical ophthalmic ointment for eye protection. The scalp was shaved and disinfected with aseptic cotton swabs. Mice were then placed in a stereotaxic frame, and a midline sagittal scalp incision (∼1 cm) was made to expose the skull, which was leveled in the bregma-lambda plane. Two stainless steel holding screws were implanted bilaterally onto the skull. A two-step procedure involving viral infection and optical fiber implantation was performed to achieve targeted gene expression and subsequent optical stimulation within the NAcSh. A microinjection needle (30-G) connected to a 10-µl Hamilton syringe was used for bilateral injections of AAV targeting the NAcSh at the following coordinates from bregma (mm): AP +1.4, ML ± 0.65, DV −4.4. The injection volume was 250 nL at a rate of 30 nL/min, and the injector was left in position for an additional 5-min period to allow for virus diffusion. Although the intended region was the NAcSh, some viruses spread occurred in the core region. Therefore, we refer to this manipulation as targeting the anterior medial NAc. After AAV injection, a zirconia ferrule with a multimode optical fiber (200 µm, Thorlabs) was implanted into the NAcSh at the following coordinates from bregma (mm): AP +1.4, ML ± 1.35, DV −3.9, with a 10° angle (Stoelting stereotaxic instrument Model Number 51730). Mice were allowed to recover for 3 weeks to obtain stable expression of ChR2, ArchT, EYFP, or tdTomato. In the D2^tdTomato^ control group, stimulation parameters were pseudo-randomly selected to match the protocols used for the D2^ChR2^ and D2^ArchT^ groups, ensuring that control mice were exposed to similar conditions without active modulation of D2 neuron activity.

### Optogenetic parameters

Mice expressing ChR2, ArchT opsins, or tdTomato were stimulated with a diode-pumped solid-state laser system with a light output intensity of 10 mW at the tip of the optical fiber patch cord, as measured with an optical power meter (PM20A, Thorlabs). For the D2^ChR2^ group, the laser output wavelength was 473 nm (OEM Laser) with a pulse duration of 10 ms delivered at 20 Hz (unless otherwise indicated for each behavioral paradigm). For the D2^ArchT^ group, the laser output wavelength was 532 nm (Laserglow Technologies), and continuous light pulses were delivered (unless otherwise indicated). The tdTomato-expressing mice received random stimulation parameters from either the ChR2 or ArchT protocols.

### Gustatory stimuli

Liquid solutions: Reagent-grade sucrose (Sigma-Aldrich) was diluted in distilled water to yield 3%, 10%, and 18% w/v sucrose solutions. Solutions were prepared fresh every other day, stored overnight at 4°C, and brought to room temperature (∼23°C) before use. Solid diets: For solid food consumption experiments, two types of food were provided: a standard chow diet (PicoLab Rodent Diet 20) and a HFD with 45% kcal fat (Research Diets).

### Behavioral tasks

All behavioral tasks were automated using an operant conditioning controller (Med Associates Inc., VT, USA). Lick events were recorded with a contact lickometer (Med Associates Inc., VT, USA). Mice were water-restricted for 23 h before the session and given access to water for 1 h afterward for tasks involving licking behavior. For food consumption tasks, mice could eat only a standard chow diet corresponding to 10% of their body weight (2-3 g) between sessions. For tasks requiring training, mice were habituated to the patch cord during training sessions by connecting them to the patch cord without laser stimulation.

### Two-bottle free-licking closed-loop assay

Mice were habituated to the behavioral box (43 x 23 cm) for two days before training. The training session aimed to familiarize mice with the two-bottle setup and establish similar licking behavior from both bottles. The box was equipped with two water bottles, with gravity-fed sippers, and connected lickometers on each side of the frontal wall. The entire test lasted six days, with one 20-min session per day. The positions of the water and sucrose bottles were randomized and counterbalanced between sessions across subjects. The procedures implemented each day were: a) Baseline (Days 1-2): Mice freely licked from both bottles without optogenetic stimulation to establish baseline sucrose preference. b) Laser Sucrose (Days 3-4): Licking the sucrose bottle triggered optogenetic stimulation. c) Laser Water (Days 5-6): Licking the water bottle triggered optogenetic stimulation. Optogenetic stimulation was delivered in a 4-s-on and 2-s-off cycle as long as the mice continued licking. The sucrose preference score was calculated as the ratio of licks received by the sucrose bottle divided by the total licks from both bottles for each day.

### Sucrose freely licking with optogenetic frequency scan

On the day before the experiment, mice were habituated to a behavioral box (22 x 18 cm, Med Associates). The task comprised five consecutive daily 20 min sessions. Each trial included three epochs: Before (laser off): The mouse had to complete three licks to initiate a new trial and enter the laser epoch. Laser epoch: On the fourth lick, the laser was activated for 1 s at different stimulation frequencies, presented in pseudo-random order: no stimulation (control) or 10 ms pulse width at 5, 8, 15, or 20 Hz. This ensured that the animal could not predict the stimulation frequency. Time out (laser off): During this 1-s epoch following laser stimulation, the laser could not be reactivated. The next trial began after the mouse completed three additional licks following the time-out epoch. Each lick was rewarded with a drop of sucrose solution (10% w/v, ∼2 μL) from a licking spout throughout the task, and the frequency-dependent effects of optogenetic stimulation on licking behavior were analyzed.

### Brief access taste test with different optogenetic stimulation frequencies

Building upon earlier investigations, this variation of the brief access task examined the effect of different stimulation frequencies (during the reward window) on motivated licking behavior. The same behavioral box (22 x 18 cm, Med Associates) was used. The task consisted of five consecutive daily 20 min sessions, each involving trials initiated by a lick on the sipper. Following the lick, a 4-s reward epoch began, signaled by an auditory cue. Every lick triggered the delivery of 10% w/v sucrose solution during this reward window. Simultaneously, with sucrose reward delivery, a laser was activated at different frequencies in a pseudorandom order: no stimulation (control), 10 ms pulse width lick-paired, 8 Hz, or 20 Hz. After the reward window ended, subsequent licks were not rewarded. To initiate a new trial, mice had to stop licking for a randomly chosen inter-trial interval (ITI) between 1 and 3 s.

### Brief access test with varying delays in optogenetic stimulation

This variation of the brief access task assessed the effect of the timing of lick-paired optogenetic stimulation on behavior. The overall structure was like the previous brief access task, with the key difference being the timing of the laser stimulation. Simultaneously with sucrose reward delivery, a laser was activated at different delay times in a pseudo-random order: no stimulation (control) or a 10 ms pulse delivered at specific delays after the lick (lick-evoked), with no delay (0 ms), a 50 ms delay, or a 100 ms delay.

### Real-time place preference (RTPP)

Mice were placed in a rectangular acrylic arena (42 x 20 x 20 cm) divided into two halves differentiated by context (visual cues: one side with black/white stripes, the other with black/white circles; Fig 5a). Mice freely explored both sides of the chamber. The experiment included two 10-min sessions separated by approximately 5 h on the same day. During each session, one side of the chamber was paired with optogenetic stimulation. Each time mice crossed to the paired chamber; they received 4 s of optogenetic stimulation followed by 2 s without stimulation. This cycle (4 s on, 2 s off) could repeat as long as the mouse remained on the paired side. The stimulation parameters were: D2^tdTomato^ and D2^ChR2^, 10 ms pulse at 20 Hz and 8 Hz stimulation (each frequency used on a separate day); D2^ArchT^ continuous pulse stimulation. The preference index for the conditioned and unconditioned sides was calculated by dividing the total time spent on each side by the total exploration time of the session. The chamber side paired with optogenetic stimulation was counterbalanced across subjects.

### Open-loop palatable (HFD) consumption test

This task investigated how mice respond to HFD consumption without context conditioning elements. Mice were placed in a white arena with the same dimensions as in the RTPP task, with an HFD pellet on each side (pellet weight measured before and after). Mice freely explored for a single 20-min session, receiving stimulation: 4 s on, 2 s off (D2^tdTomato^ and D2^ChR2^, 20 Hz, 10 ms pulse; D2^ArchT^, Continuous pulse). Using the same stimulation parameters as RTPP isolated the effects of optogenetic stimulation on HFD consumption from context conditioning. Food consumption was calculated by dividing the pellet weight difference by session time.

### RTPP with palatable (HFD) consumption

Similar to the RTPP task, mice were placed in a rectangular acrylic arena (same dimensions) with an HFD pellet placed on each side. The pellet weight was determined before and after each session. Mice freely explored both sides. The experiment consisted of five consecutive daily sessions: a) Pre-Test (Day 1, 10 min): Mice explored with HFD pellets but no optogenetic stimulation; b) Acquisition (Days 2-4, 20 min): Based on the pre-test exploration time, mice were assigned a conditioned side (more Time for D2^tdTomato^ and D2^ArchT^ or less preferred for D2^ChR2^). When crossing to and remaining on the conditioned side, they received optogenetic stimulation: D2^tdTomato^ and D2^ChR2^: 4 s on (20 Hz, 10 ms pulse width), 2 s off; D2^ArchT^: Continuous pulse; c) Test (Day 5, 10 min): Mice explored with HFD pellets without stimulation. The preference index was calculated by dividing the time spent on each side by the total exploration time. Food consumption was calculated by dividing the pellet weight difference by session time.

### Negative reinforcement test

Mice were tested for negative reinforcement in a white open circular arena (50 cm diameter) where they could freely explore during daily 30-min sessions for 3 consecutive days. Behavior was monitored from above using a video camera, and the mouse’s position was analyzed in real-time using custom MATLAB code. The code calculated the centroid of the mouse and sent this information to the operant conditioning controller (Med Associates Inc., VT, USA). The arena was virtually divided into two zones: an inner circle (10 cm diameter) designated as the ‘off zone’ and an outer annulus surrounding the off zone defined as the ‘on zone.’ The on zone was paired with optogenetic stimulation, and when a mouse entered the on zone, it received 4 s of optogenetic stimulation followed by 2 s without stimulation. This cycle could be repeated if the mouse remained in the on zone. The stimulation parameters were D2^tdTomato^ and D2^ChR2^, 10 ms pulse at 20 Hz; D2^ArchT^, optogenetic stimulation was delivered as continuous pulses. To stop optogenetic stimulation, the mouse had to enter the off zone and remain there. The analysis software determined zone entry and exit based on the position of the calculated centroid provided by the MATLAB code. The percentage of time spent in the stimulated and unstimulated zones was calculated by dividing the total time spent in each zone by the total exploration time of the session.

### Histology

After the experimental sessions, mice received a pentobarbital sodium overdose and were transcardially perfused with a phosphate-buffered saline solution followed by 4% paraformaldehyde (PFA). Brains were removed, stored for 1 day in 4% PFA, and then transferred to 30% sucrose in phosphate-buffered saline solution. Brains were sectioned into 40-μm coronal slices and mounted in a medium (Dako) for confocal microscopy to confirm viral expression and fiber placement. Mice with incorrect expression or fiber placement were excluded from the analysis. Images were taken with a confocal microscope, and contrast was improved with Adobe Photoshop CS5.1 software.

### Quantification and statistical analysis

Data were analyzed in MATLAB R2024a (The MathWorks Inc.) and GraphPad Prism 9. Data were expressed as means ± SEM. Statistical analyses were performed by first assessing normality distribution, homoscedasticity, and sample size to decide the use of parametric or non-parametric tests. As most of the data met the assumptions for parametric tests, one-way ANOVA was used, followed by a Tukey-Kramer post hoc test with an ⍶ level of 0.05. Sample sizes were decided based on previous studies using similar behavioral paradigms and are reported in figure legends. Potential confounders, such as variations in individual animal behavior and environmental factors, were controlled by randomizing the order of experimental conditions and balancing the groups by sex and age.

## Results

### Optogenetic activation of NAc D2-expressing neurons at 20 Hz reduces sucrose preference by disrupting the microstructure of licking and increasing self-stimulation

A two-bottle freely licking preference test was conducted to assess the role of NAc D2-expressing neurons in modulating sucrose preference. This experiment was conducted under three conditions, each lasting two days: 1) Baseline (No-laser), 2) Laser activated by the sucrose bottle, and 3) Laser activated by the water bottle. In the D2^tdTomato^ (control) and D2^ArchT^ (inhibition) groups, animals maintained a stable preference for sucrose over water throughout all conditions. However, in the D2^ChR2^ (activation) group, optogenetic stimulation at 20 Hz during sucrose consumption significantly reduced sucrose preference (**Fig 1c** [* one-way ANOVA; main factor: groups (Laser sucrose condition); F_(2, 62)_ = 9.20, p < 0.001]), a *post hoc* analysis further revealed that D2^ChR2^ exhibited a significant decrease compared to D2^tdTomato^ and D2^ArchT^ (all *p’s < 0.003). This reduction in sucrose preference was accompanied by a marked increase in water preference, as indicated by fewer total sucrose licks and more water licks in the D2^ChR2^ group (**Fig 1d** [* one-way ANOVA; main factor: groups (within Laser sucrose condition); F_(1, 40)_ = 13.81, p < 0.001], [# one-way ANOVA; main factor: conditions; for water F_(1, 44)_ = 24.13, p < 0.001; for sucrose F_(1, 44)_ = 13.30, p < 0.001]). As well as an increase in total licks (water+ sucrose) within the same condition for the D2^ArchT^ group [^@^ one-way ANOVA; main factor: conditions; F_(1,39)_ = 4.17, p = 0.04]. Additionally, when stimulation was paired with the water bottle, the D2^ChR2^ group exhibited a significant increase in sucrose licking compared to the baseline condition [^#^ one-way ANOVA; main factor: conditions; F_(1,36)_ = 7.75, p = 0.008], the D2^tdTomato^ group [* one-way ANOVA; main factor: groups; F_(1,26)_ =27.91, p<0.001] and generating a significant increase for total licks [^@^ one-way ANOVA; main factor: groups; F_(1,26)_ = 31.66, p<0.001], as well the D2^ArchT^ group showed a significant increase compared with D2^tdTomato^ group [* one-way ANOVA; main factor: groups; F_(1,21)_ = 12.40, p = 0.002], resulting in a significant increase in total licks [^@^ one-way ANOVA; main factor: groups; F_(1,21)_ = 11.09, p = 0.003]. This increase in licking behavior was accompanied by a higher number of trials started (a measure of self-stimulations) (**Fig 1e** [*one-way ANOVA; main factor: groups; F_(2, 62)_ = 11.67, p < 0.001]), suggesting a rewarding effect induced by 20 Hz stimulation. Additionally, comparisons between the laser and No-laser in Baseline condition in the D2^ChR2^ group showed significant differences for sucrose licks (L_s) (paired t-test; t_(44)_ = 2.72, p = 0.0092) and water licks (L_w) (paired t-test; t_(36)_ = 3.71, p = 0.0007). A deeper analysis of the licking microstructure revealed that D2 neuron activation disrupted normal licking patterns (**S1 Table**). Heatmaps of licking behavior (**Figs 1f-h**) revealed consistent licking patterns across all groups during baseline conditions (**Fig 1f**). However, optogenetic stimulation paired with sucrose consumption (**Fig 1g**) induced a significant decrease in lick bout durations, specifically within the D2^ChR2^ group [one-way ANOVA; main factor: groups (within Laser sucrose condition) F_(1, 44)_ = 35.90, p < 0.001] and reduced bout size [one-way ANOVA F_(1, 44)_ = 13.30, p < 0.001], accompanied by increased number of licking bouts. A similar fragmentation in licking microstructure was observed when water was paired with stimulation (**Fig 1h**) for bout durations [one-way ANOVA; groups; F_(1, 36)_ = 7.73, p < 0.001] and bout size [one-way ANOVA; groups; F_(1, 36)_ = 10.10, p < 0.001]. Further analysis of licking microstructural parameters (**S1 Table**) confirmed that optogenetic activation of D2 neurons significantly reduced bout size and duration while increasing bout frequency for water in the D2^ChR2^ group. These findings indicate that optogenetic activation of NAc D2-expressing neurons disrupts normal licking behavior and induces self-stimulation, suggesting a rewarding effect. This suggests that the observed decrease in sucrose preference likely stems from the combined rewarding and lick-impairing effects of optogenetic stimulation.

**Figure 1.**
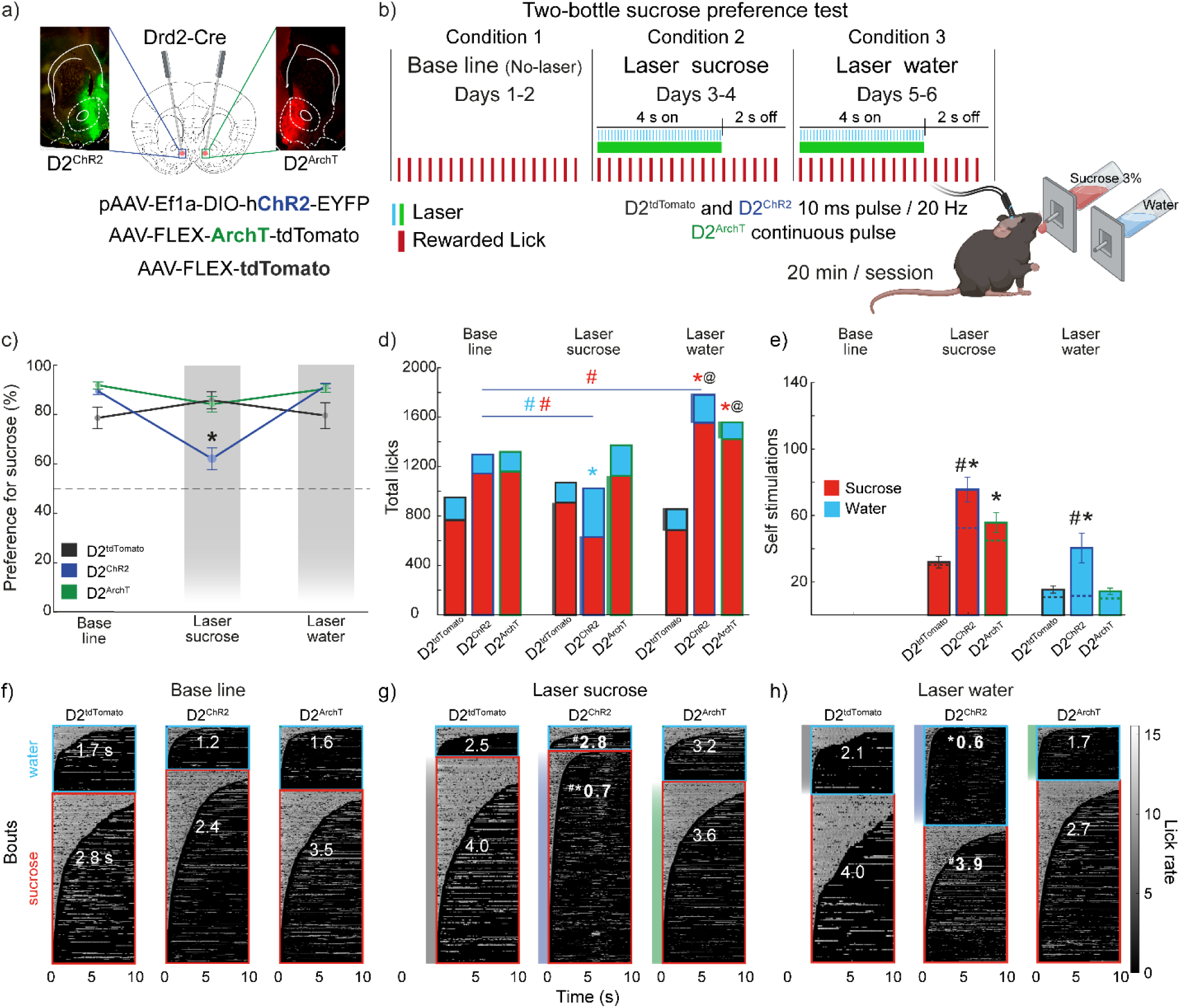
Optogenetic activation of NAc D2 neurons at 20 Hz reduces sucrose preference, disrupts licking microstructure, and increases self-stimulation trials. (a) Coronal brain sections depicting Cre recombinase expression in D2-expressing neurons. Below are the viral constructs used: pAAV-Ef1a-DIO-hChR2-EYFP (activation), AAV-FLEX-ArchT-tdTomato (inhibition), and AAV-FLEX-tdTomato (control). (b) Two-bottle test protocol: baseline (No-laser, Days 1-2), laser triggered by licking the sucrose bottle (Days 3-4), and laser triggered by the water bottle (Days 5-6). (c) Sucrose preference (%) across conditions in each group. (d) Total licks during baseline, laser sucrose, and laser water conditions. (e) Total trials (number of trials with self-activated laser). Dashed lines represent the baseline, calculated from the licks given per each solution in the off condition and the trial time (4 seconds on, 2 seconds off), which reflects the expected times the mice would start a new trial if laser stimulation were present. (f-h) Heatmaps of licking bouts. Water-licking bouts are shown in blue, and sucrose-licking bouts in red. (f) Baseline condition. (g) 4 s laser triggered with sucrose bottle. (h) Laser triggered with a water bottle. Licking rate (licks per second) is shown on the right. * significant differences with the control group (D2^tdTomato^) within the same condition; # significant differences compared to baseline within the same group. @ significant difference for total licks (water + sucrose) against control group within the same condition. All significant differences had p < 0.05. Data are mean ± SEM. The statistical analyses were conducted with one-way ANOVA followed by Tukey-Kramer post hoc tests. D2^tdTomato^ n=5, D2^ChR2^ n=11, D2^ArchT^ n=9. Licking microstructure details are provided in S1 Table.

### Optogenetic frequency-dependent modulation of licking behavior by NAc D2-expressing neurons

To assess the impact of stimulation frequency on licking behavior, we used a freely licking task with a laser scanning frequency test, as previously reported, to examine frequency-dependent activation of D2 NAc neurons.[14] The task operates in a cycle of three epochs: Before (4 licks), Laser (1 s), and Timeout (1 s). A new cycle is triggered by four consecutive licks immediately after the Timeout epoch. In this task, control animals (D2^tdTomato^) exhibited consistent lick rates across all laser frequencies (**Fig 2b**). In contrast, D2^ChR2^ activation reduced the lick rate compared to the No-laser trials, with stronger reductions observed at higher frequencies, especially 8, 15, and 20 Hz (**Figs 2c, 2f, and 2i**). Cumulative lick distributions during laser epochs revealed significant reductions in the D2^ChR2^ group across all frequencies compared to No-laser trials [^#^ one-way ANOVA; main factor: types of trials; F_(4, 625)_ = 38.94, p < 0.001, all Tukey-Kramer; p<0.001], and the control group (D2^tdTomatto)^ with the most pronounced reductions at 8 Hz [* one-way ANOVA; main factor: groups; F_(4, 354)_ = 33.98, p < 0.001], 15 Hz [* one-way ANOVA; main factor: groups; F_(4, 354)_ = 39.93, p < 0.001], and 20 Hz [* one-way ANOVA; main factor: groups; F_(4, 354)_ = 52.72, p < 0.001] (all ^#,@^ p’s < 0.001) (**Fig 2i**). Additionally, the 5 Hz stimulation allowed significantly more licks than higher frequencies (8, 15, and 20 Hz) within the D2^ChR2^ group [one-way ANOVA; 5 Hz vs. other frequencies; F_(3, 187)_ = 8.79, p < 0.001]. Inhibition of D2 neurons (D2^ArchT^) at 5 Hz stimulation generated significantly more licks compared to the control group (**Figs 2j**) [*one-way ANOVA; main factor: groups; F_(2, 354)_ = 18.69, p <0.001], while there were no significant differences at 8, 15, or 20 Hz compared to the control group (**Fig 2j**). During the Laser epoch, D2^ChR2^ animals displayed significantly fewer licks at 8, 15, and 20 Hz compared to No-laser trials [^#^one-way ANOVA; main factor: trial types; F_(4, 130)_ = 14.07, p < 0.001] (**Fig 2k**). Inhibition of D2 neurons (D2^ArchT^) at 20 Hz resulted in significant suppression of licking behavior [^#^one-way ANOVA; main factor: trial types; F_(4, 160)_ = 3.44, p = 0.009] (**Fig 2k**). Stimulation effects persisted into the post-laser timeout period, with a tendency for lick frequency to decrease with increasing stimulation frequency (**Fig 2l**). Although this change was less pronounced during the Timeout epoch, the tendency suggests a continued stimulation influence. These results suggest that NAc D2 neuron activity modulates licking behavior in a frequency-dependent manner, acting as a “brake” on consummatory actions and decreasing sucrose intake. Higher stimulation frequencies, particularly at 15 and 20 Hz, produce stronger suppression and have long-lasting effects on re-initiating reward-seeking behavior.

**Figure 2.**
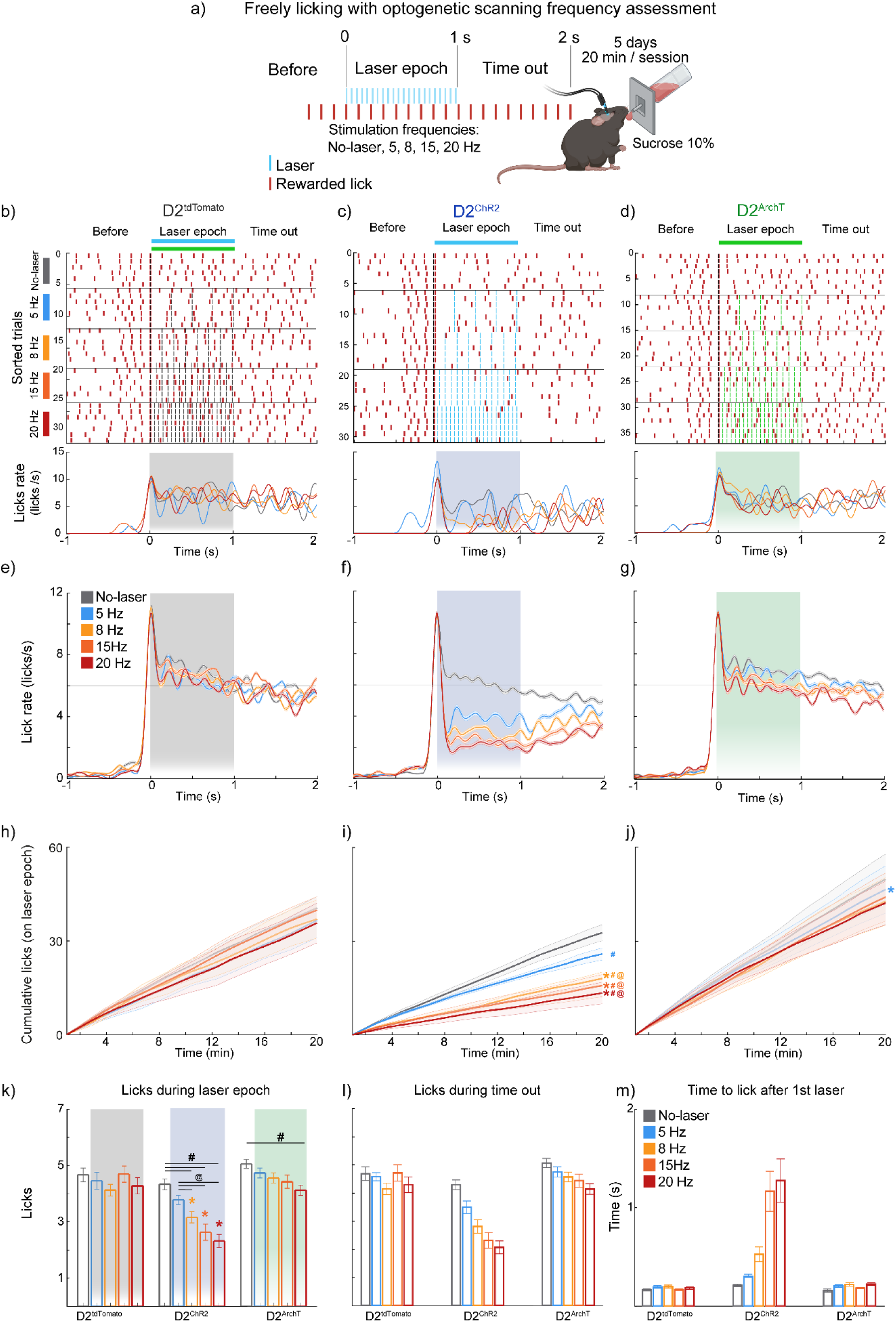
Optogenetic activation of NAc D2-expressing neurons disrupts licking behavior in a frequency-dependent manner. (**a**) Schematic representation of the freely licking test with 10% sucrose and an optogenetic scanning frequency. The freely licking task is composed of three epochs: Before (four licks), Laser (1 s), and Time out (1 s). Every four licks given after the time-out epoch starts a new Laser epoch. (**b-d**) Representative raster plots showing individual licks (red ticks) and laser pulses (blue for activation, green for inhibition) during a single session in D2^tdTomato^ (control), D2^ChR2^ (activation), and D2^ArchT^ (inhibition) groups. The figures below correspond to the Peri-Stimulus Time Histogram (PSTH) of the single session. (**e-g**) Population PSTHs display mean lick rates for all mice aligned to laser stimulation onset (Time = 0 s). Note that the licking structure is disrupted in a frequency-dependent manner. (**h-j**) Cumulative lick distributions during laser epochs across the session confirmed the disruption of licking for the D2^ChR2^ group. (**k**) Total licks during the 1 s laser epoch reveal frequency-dependent disruption in the D2^ChR2^ group. (**l**) Licks during the 1 s timeout phase (post-laser) show recovery. (**m**) The time to start licking after the first laser pulse (Time = 0 s) shows delayed re-initiation in the D2^ChR2^ group, increasing with frequency. * Significant differences from the control group (D2^tdTomato^) within the same condition; # significant differences compared to No-laser, @ significant differences compared to 5 Hz frequency within the same group. All significant differences had p < 0.05. Data are mean ± SEM. The statistical analyses were conducted with one-way ANOVA followed by Tukey-Kramer post hoc tests. D2^tdTomato^ n=3, D2^ChR2^ n=8, D2^ArchT^ n=7. Licking microstructure details are provided in **S2 Table**.

### Self-paced (unlike experimenter-imposed frequency) optogenetic stimulation of D2 neurons attenuates disruption of licking in a brief access taste test

We used the brief access taste test (BATT) to investigate whether the frequency-dependent effects of stimulation on licking behavior persisted under a self-paced lick-driven rhythmic stimulation pattern. In this task, mice could lick for a 10% sucrose (Suc) solution during a 4 s reward period with different optogenetic stimulation trials (No-laser, lick-paired (0 ms delay), 8 and 20 Hz). The D2^tdTomato^ (control) (**Figs 3b, 3e**) group maintained consistent licking behavior across all trials, showing no significant disruption by these modulations. In contrast, based on the optogenetic stimulation pattern, the D2^ChR2^ (activation) group exhibited a significant difference in licking behavior (**Figs 3c, 3f**). Notably, 20 Hz stimulation significantly reduced the licking rate [^#^one-way ANOVA; main factor; trial types F_(4, 416)_ = 22.3, p <0.001], while lick-paired stimulation preserved a licking rate similar to the No-laser condition, highlighting the importance of self-timing in optogenetic stimulation and its effects on licking behavior.

**Figure 3.**
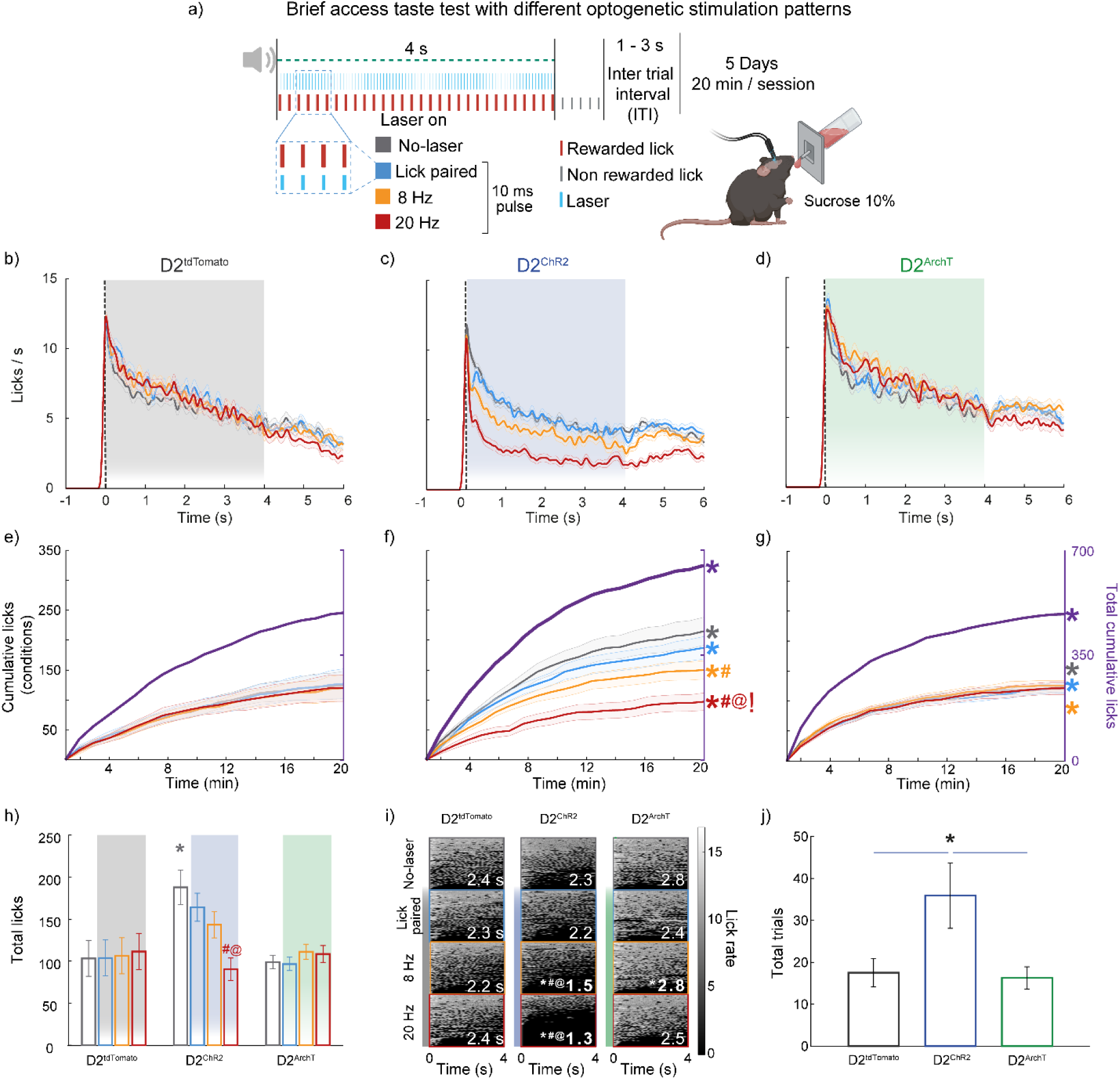
Frequency-dependent disruption of licking behavior in a sucrose brief access test demonstrates the advantage of self-paced (lick-paired) activation over imposed optogenetic stimulation and exhibits enhanced task engagement through D2 neuron activation. (**a**) Schematic representation of the sucrose brief access (4 s) taste test with optogenetic stimulation, including No-laser (gray) and 10 ms pulses at 0 ms delay (blue), 8 Hz (yellow), and 20 Hz (red) frequencies for 4 s. Each trial included the delivery of sucrose in each lick for 4 s (red ticks); after that, additional licks were empty (gray ticks). To start a new trial, mice must stop licking for a 1-3 s inter-trial interval (ITI). The experiment comprises 5 sessions, each lasting 20 min. (**b-d**) PSTH displaying mean sucrose lick rates during the 6 s of trial for all D2^tdTomato^ (**b**), D2^ChR2^ (**c**), and D2^ArchT^ mice **(d**). The shaded area represents the 4 s laser stimulation period. (**e-g**) Cumulative licks over the 20-minute session for all stimulation patterns within the same group are shown on the left y-axis, with total cumulative licks (in purple) on the right y-axis. (**h**) Total licks per session for each group under different stimulation conditions. (**i**) Heatmaps plotting licking bouts across stimulation patterns for all groups, with the x-axis representing time in s (0-4 s) from lick bout onset and the y-axis showing individual bouts (the number in white indicates lick bout duration in s). Dark gray edges (bar line at left) represent No-laser bouts, blue edges lick-paired laser stimulation, yellow edges 8 Hz imposed stimulation, and red edges 20 Hz stimulation. Light gray intensity indicates lick rate. (**j**) Total trials given by each group under different stimulation conditions. The activation of D2 neurons in the NAc led to significantly more trials started in the D2^ChR2^ group than in the D2^tdTomato^ group. Colored symbols: * significant difference compared to D2^tdTomato^ at the same frequency, # significant difference compared to the No-laser condition within the same group, and @ significant difference between the lick-paired condition and the 8 Hz or 20 Hz conditions within the same group. The symbol ! indicates a significant difference between the 8 Hz and 20 Hz conditions within the same group. All significant differences had p < 0.05. Data are mean ± SEM. The statistical analyses were conducted with one-way ANOVA followed by Tukey-Kramer post hoc tests. D2^tdTomato^ n=6, D2^ChR2^ n=5, D2^ArchT^ n=5. Further analysis of the licking microstructure is presented in **S3 Table**.

These effects were also reflected in cumulative licks across the session, where the D2^ChR2^ group exhibited significantly fewer licks during stimulation at 20 Hz than the other trial types (**Fig 3f** [^#^one-way ANOVA; main factor: trial types; F_(3, 416)_ = 22.63, p < 0.001]). A *post hoc* analysis confirmed that the No-laser and lick-paired conditions significantly differed in the D2^ChR2^ group across trial types [^#^p < 0.001], underscoring the impact of stimulation frequency on licking behavior. The 8 Hz imposed stimulation also reduced cumulative licks compared to the No-laser condition [p<0.001], although this reduction was less pronounced than that induced by the 20 Hz stimulation [^!^p<0.001]. While the D2^ArchT^ (inhibition) group significantly increased the cumulative licks in the total licks and in the lick paired, 8, 20 Hz trials (**Figs 3d, 3g**) compared to the control group (all *p’s<0.01) (**Figures 3e, 3g**).

We plotted the total licks per session to observe the overall impact on licking behavior (**Fig 3h**). This plot further emphasized these differences across groups [*one-way ANOVA; main factor: groups F_(2, 65)_ = 7.78, p <0.001]. In the D2^ChR2^ group [*one-way ANOVA; main factor: trial types F_(3, 80)_ = 7.668, p <0.001], total licks during the 20 Hz condition were significantly reduced compared to the No-laser(^#^p <0.001) and lick-paired conditions(^@^p <0.001)). Conversely, total licks in the D2^tdTomato^ and D2^ArchT^ groups showed no significant differences across stimulation types, confirming that optogenetic activation of D2-expressing neurons, rather than inhibition, more strongly affected licking behavior.

Importantly, the increased licking behavior observed during control trials in the D2^ChR2^ group may be explained by the sequential nature of the task. Given that each trial must be completed before passing to the next one, the animals could display an enhanced motivation effect from previous trials where the D2 neurons were activated (panel **h**). Microstructural analysis of licking behavior (**Fig 3i)** revealed, at 8 [*one-way ANOVA; main factor: groups; F_(2, 65)_ = 12.50, p <0.001] and 20 Hz [*one-way ANOVA; main factor: groups; F_(2, 65)_ = 10.03, p <0.001] frequencies, more fragmented licking in the D2^ChR2^ group characterized by shorter bout duration and increased bout number [*one-way ANOVA; group; F_(2, 65)_ = 16.95, p < 0.001]. Interestingly, despite the disrupted licking behavior, the D2^ChR2^ group initiated more trials than the other groups (**Fig 3j** [*one-way ANOVA; groups; F_(2, 65)_ = 14.14, p < 0.001]), indicating that 20 Hz stimulation disrupted consummatory licking but increased the persistence in the task. **S3 Table** shows that high-frequency stimulation (20 Hz) significantly reduced lick rate, bout size, and bout duration in the D2^ChR2^ group while increasing the number of licking bouts. This suggests that while high-frequency stimulation disrupts the typical licking behavior, it paradoxically enhances the animal’s incentive motivation to persist in seeking the reward, as evidenced by increased trial initiation.

### Lick-paired optogenetic activation with delayed timing of NAc D2 neurons slightly disrupts the licking microstructure but enhances sucrose intake

Given the advantage of lick-paired over imposed stimulation, we next assessed the effect of lick-paired but using multiple delays relative to the lick tongue protrusion-retraction cycle. To this aim, we used a variation of the brief access test, where optogenetic activation was applied with different delays relative to the tongue contacting the sipper (0, 50, and 100 ms) after each lick, alongside no-laser trials that served as controls. Licking rates remained relatively stable along the reward epoch across all trial types and groups (**Figs 4b-d**), indicating that delayed activation or inhibition had minimal effects on the overall licking behavior. However, despite these stable rates, the D2^ChR2^ group exhibited a significantly higher total number of cumulative licks than the D2^tdTomato^ group across all trial types (**Figs. 4e-f** [*one-way ANOVA; main group: No-laser F_(2, 312)_ = 11.05; p < 0.001; main group: 0 ms F_(2, 312)_ = 3.29, p = 0.04; main group: 50 ms F_(2, 312)_ = 12.49, p <0.001; main group: 100 ms F_(2, 312)_ = 18.67, p <0.001; main group: Total cumulative licks F_(2, 1257)_ = 40.15, p <0.001]. The D2^ArchT^ group also showed significant increases in total cumulative licks (*p = 0.002), particularly in the 100 ms delay trials (*p < 0.001). These findings indicate a significant increase in the total number of cumulative licks for the D2^ChR2^ group, suggesting greater overall licking activity than the other groups.

**Figure 4.**
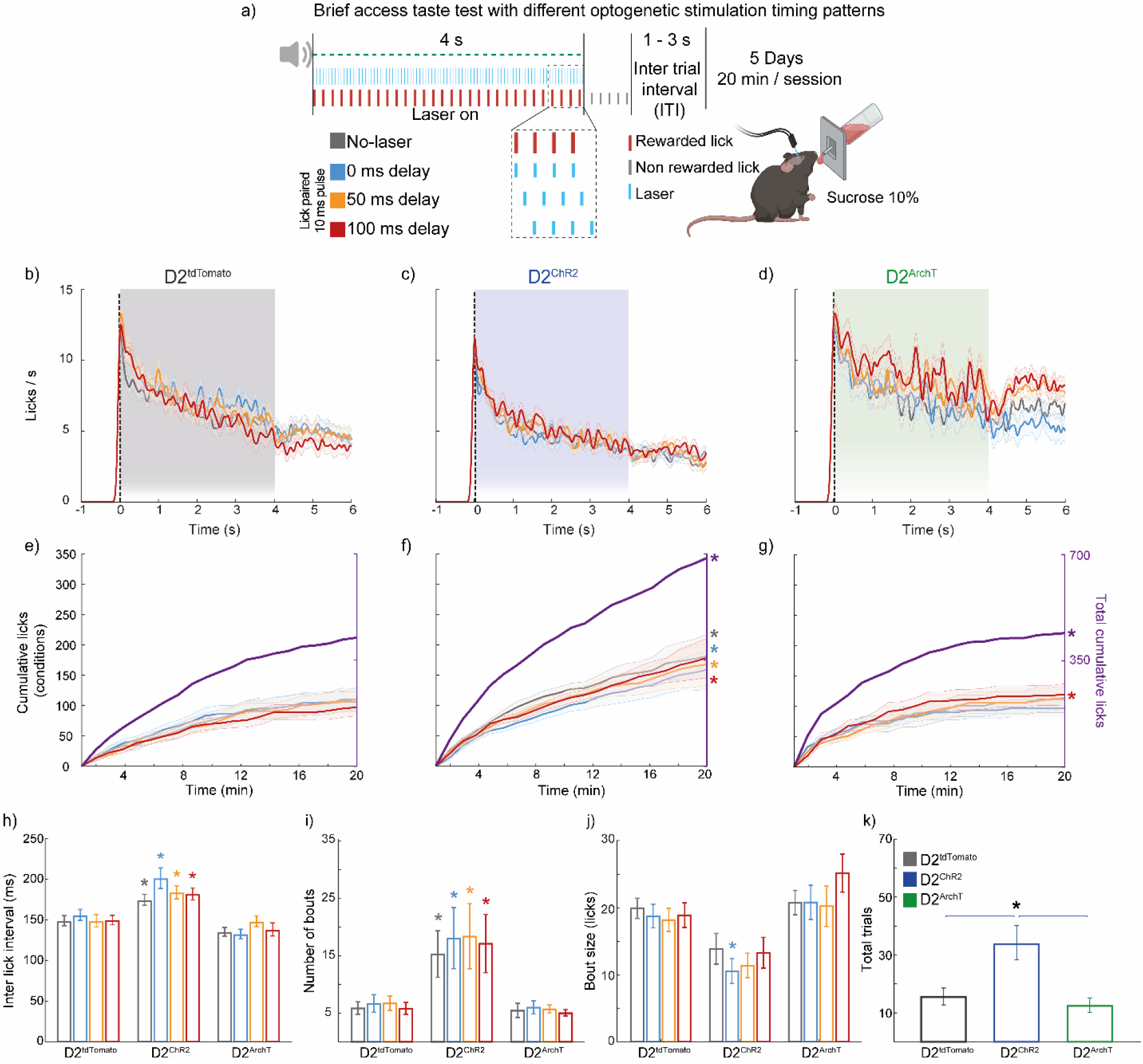
Lick-paired optogenetic stimulation of D2 NAc neuron fragments licking microstructure but paradoxically promotes sucrose intake through its rewarding effects. (**a**) Schematic representation of the brief access test with optogenetic stimulation. Mice were subjected to varying delays (0, 50, and 100 ms) of 10 ms pulses of laser stimulation entrained to licking during a 4 s reward period alongside a No-laser condition. (**b-d**) PSTHs display the mean sucrose lick rates during the 4 s laser stimulation period for D2^tdTomato^ (**b**), D2^ChR2^ (**c**), and D2^ArchT^ (**d**). (**e-g**) Cumulative licks over the 20 min session are shown for all conditions within the same group on the left y-axis, while total cumulative licks (in purple) are shown on the right y-axis. (**h**) Inter-lick interval (ms) for each group under different stimulation conditions. In the D2^ChR2^ group, inter-lick intervals across all conditions were significantly longer compared to the D2^tdTomato^ group in the same condition. In the D2^ArchT^ group, the 0 ms delay condition showed a significantly shorter inter-lick interval than the D2^tdTomato^ group. (**i**) Number of licking bouts for each group across conditions. The D2^ChR2^ group exhibited significantly more bouts than the D2^tdTomato^ group across all frequencies. (**j**) Bout size (licks) for each group. The D2^ChR2^ group had significantly smaller bout sizes than the D2^tdTomato^ group in the No-laser, 0 ms delay, and 50 ms delay conditions. (**k**) Total trials initiated by each group under different stimulation conditions. The activation of D2 neurons in the NAc led to significantly more trials initiated in the D2^ChR2^ group than in the D2^tdTomato^ control group. Colored symbols: * significant difference compared to D2^tdTomato^ (control) at the same frequency, significant difference compared to the No-laser condition within the same group. All significant differences had p < 0.05. Data are mean ± SEM. The statistical analyses were conducted with one-way ANOVA followed by Tukey-Kramer post hoc tests. D2^tdTomato^ n=6, D2^ChR2^ n=5, and D2^ArchT^ n=5. Further analysis of the licking microstructure is presented in **S4 Table**.

Although overall, the lick-paired protocol was phased-locked to the ‘natural’ licking rhythm than imposed frequency stimulation, we found that the inter-lick interval was significantly longer in the D2^ChR2^ compared to the D2^tdTomato^ group across all conditions (**Fig 4h** [*one-way ANOVA; main factor: group: No-laser F_(2, 30)_ = 8.35, p = 0.01; 0 ms F_(2, 30)_ = 13.95, p < 0.001; 50 ms F_(2, 30)_ = 7.57, p = 0.002; 100 ms F_(2, 30)_ = 13.55, p <0.001]). This behavioral protocol also revealed the disruptive effect of optogenetic activation on rhythmic oromotor components of licking behavior, even with lick-paired stimulation patterns.

Regarding licking bouts, the D2^ChR2^ group showed a significantly higher number of bouts than the D2^tdTomato^ group across all conditions (**Fig 4i** [*all p’s < 0.001]). However, the size of each lick bout was smaller in the D2^ChR2^ group compared to the D2^tdTomato^ group under the 0 ms delay condition (**Fig 4j** [*one-way ANOVA; main factor: group F_(2,30)_ = 6.52, p = 0.01]).

Finally, despite the evident fragmentation of licking behavior, the D2^ChR2^ group initiated significantly more trials compared to both the D2^tdTomato^ and D2^ArchT^ groups (**Fig 4k** [one-way ANOVA; main factor: group; F_(2, 30)_ = 15.41, p < 0.001]), underscoring a paradoxical increase in task engagement and reward driven by D2 neuron activation. These findings indicate that although self-paced rhythmic optogenetic activation of NAc D2-expressing neurons affects licking microstructure, this effect is less pronounced than when the experimenter imposes the stimulation frequency. Interestingly, this self-paced activation paradoxically increases motivation and task persistence, leading to greater sucrose intake, especially when the activation is paired with physiological licking rhythms.

### Optogenetic activation of NAc D2-expressing neurons at 20 Hz (but not 8 Hz) drives place preference and enhances exploratory behavior

Next, we explored how optogenetic activation of NAc D2-expressing neurons modulates spatial contextual rewards beyond licking behavior. We assessed the effects of different stimulation frequencies on place preference using a real-time place preference (RTPP) assay. Optogenetic stimulation was delivered at 8 or 20 Hz in D2^ChR2^ and D2^tdTomato^ groups in this paradigm.

Heatmaps illustrating the time spent in a spatial location (in yellow; **Figs 5b-e**) revealed a clear preference for the stimulated side in the D2^ChR2^ group when stimulated at 20 Hz (**Fig 5c**). Mice in this group spent significantly more time on the stimulated side during both sessions compared to the D2^tdTomato^ control and the 8 Hz stimulation condition (**Fig 5f** [*one-way ANOVA; main factor: group; F_(3, 22)_ = 7.19, p < 0.001]). In contrast, no significant place preference was observed in the D2^tdTomato^, D2^ChR2^ 8 Hz, and D2^tdTomato^ 20 Hz groups.

**Figure 5.**
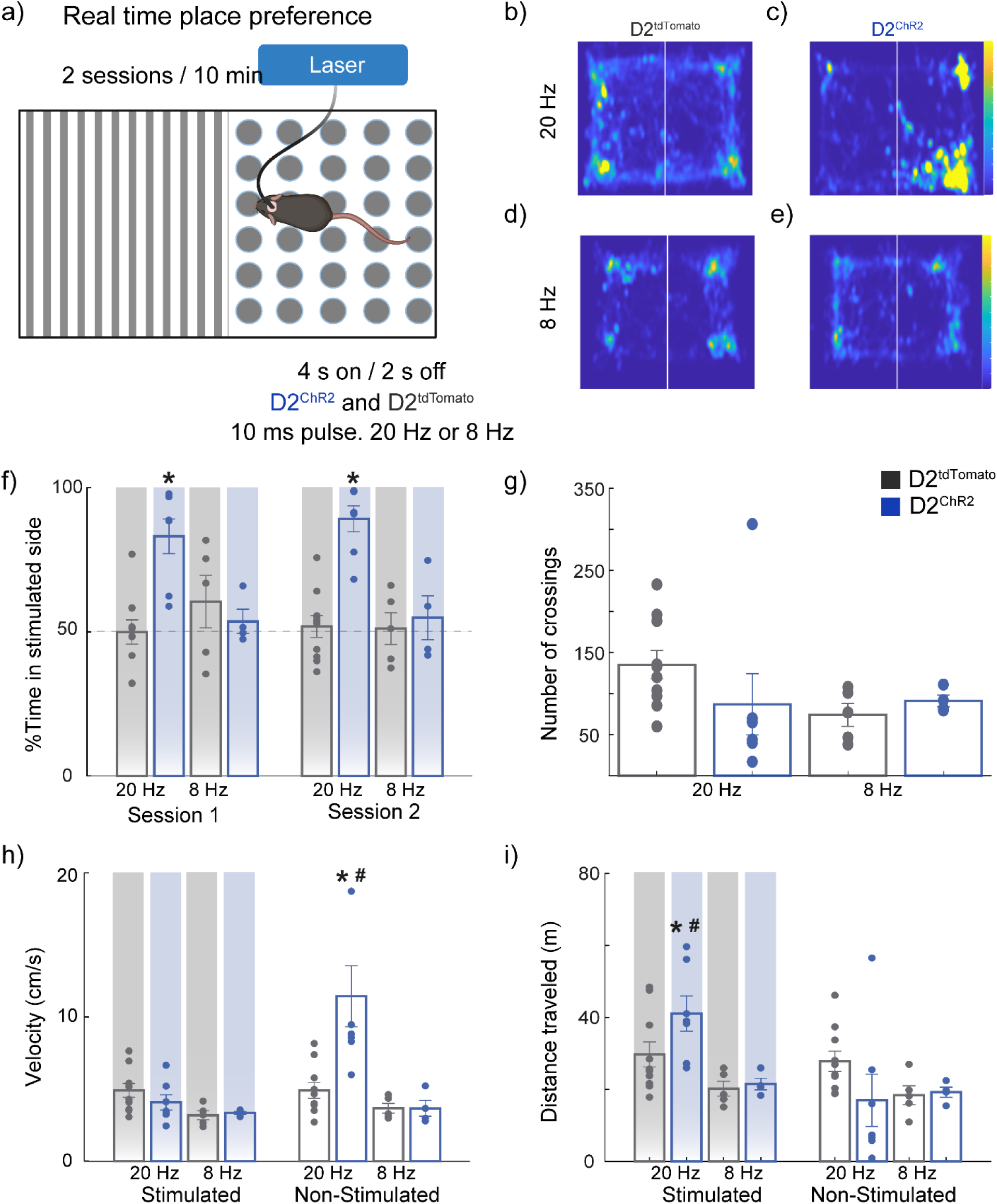
Optogenetic activation of NAc D2 neurons at 20 Hz (but not 8 Hz) is rewarding and consistently drives place preference. (**a**) Schematic representation of the Real-Time Place Preference (RTPP) task: Mice explored a rectangular acrylic chamber divided into two visually distinct halves (one with black/white stripes and the other with black/white circles). Upon entering the paired side, mice received optogenetic stimulation for 4 s(10 ms pulse width at 20 Hz or 8 Hz for D2^tdTomato^ and D2^ChR2^), followed by 2 s off, as long as mice remained on that side. Mice performed two 10 min sessions, separated by approximately 5 h. The side paired with stimulation was counterbalanced across subjects. (**b-e**) Heatmaps of mouse locations during the task under different conditions: (**b**) D2^tdTomato^ with 20 Hz stimulation, (**c**) D2^ChR2^ with 20 Hz stimulation, (**d**) D2^tdTomato^ with 8 Hz stimulation, and (**e**) D2^ChR2^ with 8 Hz stimulation (**f**) Percentage of time spent on the stimulated side during Session 1 and Session 2. Mice expressing D2^ChR2^ at 20 Hz spent significantly more time on the stimulated side compared to D2^tdTomato^ controls and the 8 Hz condition. No significant preference was observed for 8 Hz groups. (**g**) Number of side crossings between the two halves of the chamber. No significant differences were found across conditions. (**h**) Velocity of the mice across conditions. The D2^ChR2^ 20 Hz group had significantly higher velocity on the non-stimulated side compared to the stimulated side and all other groups and conditions on the non-stimulated side. (**i**) Total distance traveled during stimulation sessions. The D2^ChR2^ 20 Hz group traveled significantly more on the stimulated side than D2^tdTomato^ controls and other conditions. * Significant difference compared to other groups in the same session or side, # significant difference in the other side within the same group. All significant differences had p < 0.05. Data are mean ± SEM. The statistical analyses were conducted with one-way ANOVA followed by Tukey-Kramer post hoc tests. D2^tdTomato^ 8 Hz n=5, D2^ChR2^ 8 Hz n=4, D2^tdTomato^ 20 Hz n=9 and D2^ChR2^ 20 Hz n=7.

The number of transitions between the two sides of the chamber did not differ significantly between groups (**Fig 5g**). However, the D2^ChR2^ 20 Hz group exhibited significantly higher velocity on the non-stimulated side compared to other groups(**Fig 5h** [*one-way ANOVA; main factor: group; F_(3, 22)_ = 8.85, p < 0.001]), and conditions (^#^p =0.01) while the total distance traveled was greater on the stimulated side (**Fig 5i** [# t-test (paired); t(6) = 3.26, p = 0.01]) and against other groups [*one-way ANOVA; main factor: group, F_(3, 22)_ = 5.36, p = 0.006]. These findings indicate that the increased locomotion was specific to the stimulated chamber, suggesting that the 20 Hz stimulation enhanced both place preference and movement within the stimulated area. These results highlight that 20 Hz stimulation of D2-expressing neurons in the NAc induces a robust rewarding effect, as evidenced by place preference, and modulates locomotor behavior, leading to increased movement within the stimulated environment.

### NAc D2 neuron inhibition induces a negative hedonic state (but not aversion) that is unmasked by an HFD choice paradigm

To investigate the role of NAc D2-expressing neurons in context-dependent behaviors, we first examined their influence on place preference using the RTPP task. Mice freely explored a chamber divided into two distinct halves, with one side paired with optogenetic stimulation. In this task, optogenetic inhibition (continuous stimulation) was delivered to D2^ArchT^ mice, while D2^tdTomato^ controls received 20 Hz stimulation (**Fig 6a**, left; D2^tdTomato^ data reused in **Fig 5**). Heatmaps representing animal spatial exploration showed no significant preference for the stimulated side in either group (**Fig 6a**, middle). Quantitatively, the percentage of time spent on the stimulated side was similar across both sessions, with no significant differences between groups (**Fig 6a**, right). These data suggest that silencing NAc D2-expressing neurons generated, *per se,* neither aversion nor preference under this standard RTPP protocol.

**Figure 6.**
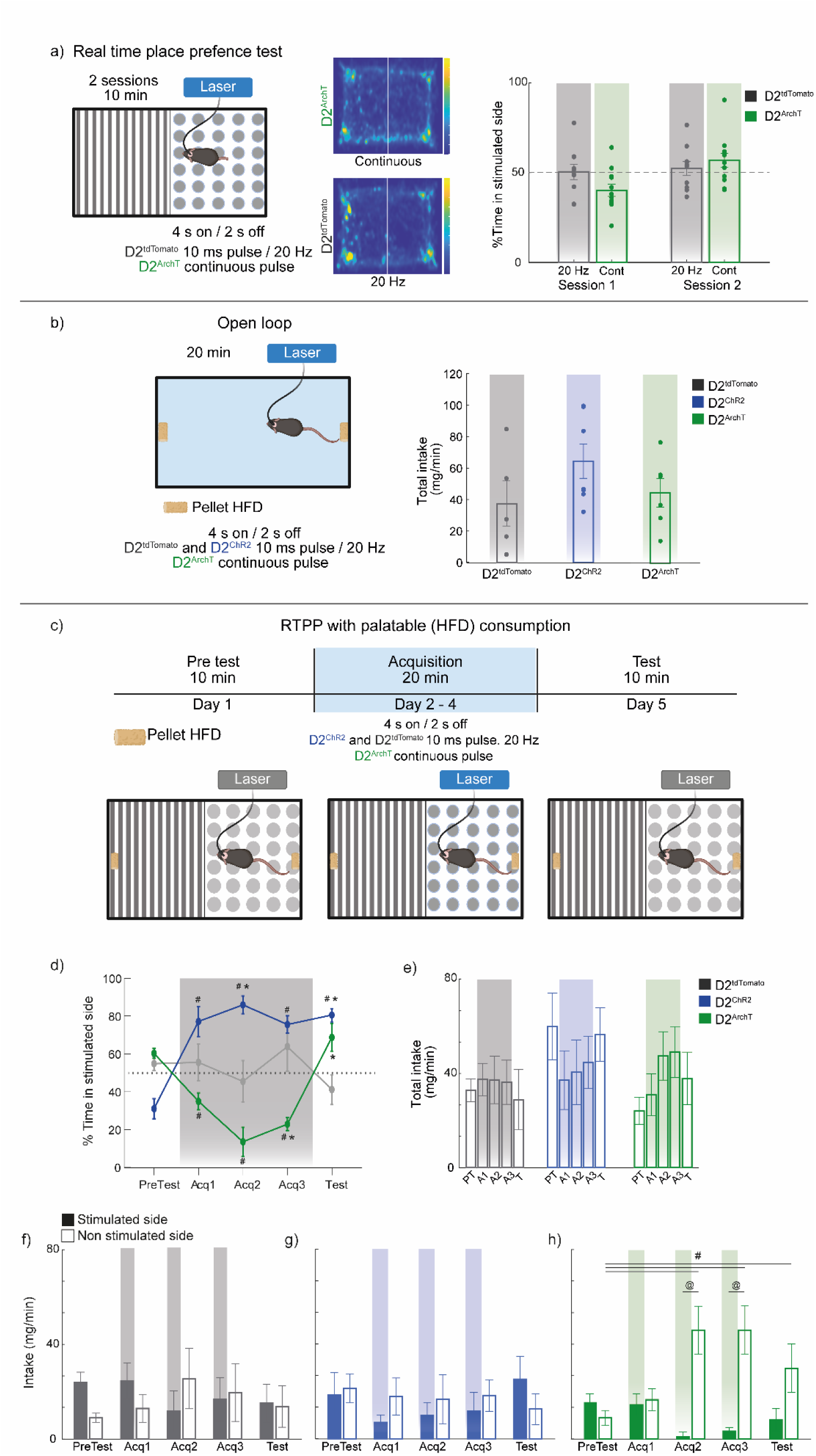
Context-dependent effects of silencing NAc D2 neurons: inhibition only alters HFD consumption based on environmental context. (**a**) Left panel. Schematic representation of the RTPP task: Mice explored a rectangular acrylic chamber divided into two visually distinct sides. Upon entering the laser paired side, mice received optogenetic stimulation for 4 s (10 ms pulses at 20 Hz for D2^tdTomato^ or continuous stimulation for D2^ArchT^), followed by 2 s off. Mice experienced two 10 min sessions, separated by approximately 5 h. The side paired with stimulation was counterbalanced across subjects. Middle panels Heatmaps of mouse locations during the task under different conditions: (middle-top) D2^ArchT^ with continuous stimulation, (middle-bottom) D2^tdTomato^ with 20 Hz stimulation. Right panel percentage of time spent on the stimulated side during session 1 and session 2. No significant preference was observed. D2^tdTomato^ 20 Hz n=9 (same data from Fig 5) and D2^ArchT^ n=12. (**b**) Left panel. Schematic of the open-loop paradigm used to assess the effects of optogenetic manipulation of D2-NAc neurons on HFD intake. optogenetic stimulation was delivered in 4 s on / 2 s off intervals, with continuous stimulation applied to the D2^ArchT^ group, while a 20 Hz frequency was used for the D2^ChR2^ and D2^tdTomato^ groups. Right panel. Total HFD intake (mg/min) for D2^tdTomato^ (control), D2^ChR2^ (activation), and D2^ArchT^ (inhibition) groups during the 20 min session. D2^tdTomato^ n=5, D2^ChR2^ n=6, D2^ArchT^ n=4. (**c**) Schematic representation of the RTPP paradigm associated with palatable food intake. Mice were conditioned to associate one side of the chamber with optogenetic stimulation (4 s on, 2 s off). D2^ChR2^ and D2^tdTomato^ groups received 10 ms pulses of 20 Hz stimulation, while the D2^ArchT^ group received continuous pulse stimulation. (**d**) Time course of the percentage of time spent on the stimulation-paired side during pre-conditioning (PreTest), conditioning (Acq1, Acq2, Acq3), and post-conditioning test (Test). Note that the D2^ChR2^ group significantly preferred the stimulated side during conditioning, consistently promoting place preference in the test without stimulation. In contrast, silencing D2 neurons induces a tendency to avoid the laser-paired side (**e**) Total HFD intake across conditioning days for all groups. There is a tendency for reduced intake during the activation of NAc D2 neurons (D2^ChR2^) and an opposite tendency for increased intake during inhibition (D2^ArchT^). (**f**-**h**) Comparison of HFD intake on the stimulated versus non-stimulated sides for D2^tdTomato^ (**f**), D2^ChR2^ (**g**), and D2^ArchT^ (**h**) groups. The D2^ArchT^ group showed a significant increase in HFD consumption on the non-stimulated side during the last two conditioning days compared to pre-conditioning. Gradient rectangles depict the condition or side paired with optogenetic stimulation. D2^tdTomato^ n=7, D2^ChR2^ n=7, D2^ArchT^ n=5: * significant difference compared to D2^tdTomato^ in the same session, # significant difference compared to the PreTest session within the same group, and @ significant difference between the stimulated and non-simulated side within the same group. All significant differences had p < 0.05. Data are mean ± SEM. The statistical analyses were conducted with one-way ANOVA followed by Tukey-Kramer post hoc tests.

Next, we assessed the effects of D2 neuron modulation on feeding behavior using an open-loop paradigm and HFD alone in one spatial context. In this protocol, mice were placed in an arena with HFD pellets on both opposite sides, while optogenetic stimulation was applied in 4 s on, 2 s off intervals for D2^tdTomato^ and D2^ChR2^ (20 Hz, 10 ms pulse width) and continuous pulse stimulation for D2^ArchT^ (**Fig 6b**, left). In this context, total HFD intake did not significantly differ between the control (D2^tdTomato^), activation (D2^ChR2^), and inhibition (D2^ArchT^) groups (**Fig 6b**, right), indicating that D2 neuron activation or inhibition *per se* was not sufficient to influence HFD feeding behavior in a neutral environment.

Finally, in a variation of the RTPP task, we explored whether NAc D2-expressing neurons modulate place preference and HFD consumption in a conditioned context, where one side of the chamber was paired with optogenetic stimulation, and the mice could choose whether or not to eat HFD pellets on the stimulated side on the box (**Fig 6c**). During conditioning, D2^ChR2^ mice developed a significant preference for the stimulation-paired side, which persisted into the post-conditioning test phase without laser (**Fig 6d** [all *p’s <0.006, see **S5 Table** for statistical analysis]), suggesting the formation of a long-term spatial reward memory. Conversely, D2^ArchT^ mice initially avoided the stimulation-paired side during acquisition [^#^one-way ANOVA; main factor: sessions; F_(4, 20)_ = 18.78, p < 0.02], but this behavior did not persist in the test phase (**Fig 6d**).

Analysis of total HFD intake revealed a tendency for reduced intake in the D2^ChR2^ group during activation, while the D2^ArchT^ group exhibited increased consumption during inhibition (**Fig 6e**). In particular, the D2^tdTomato^ (**Fig 6f**) and D2^ChR2^ (**Fig 6g**) groups had no change in the HFD intake between the stimulated and non-stimulated side, while D2^ArchT^ mice significantly increased HFD intake on the non-stimulated side during the final acquisition days, an effect that persisted into the test phase (**Fig 6h** [^#^one-way ANOVA; main factor: side; F_(4,25)_ = 4.14, p = 0.01] [t-test; Adq2 t_(5)_ = −4.40, p = 0.007; Adq3 t_(5)_ = - 4.39, p = 0.007]). This suggests that inhibiting NAc D2-expressing neurons induces negative hedonics, increasing palatable food consumption on the non-stimulated side. These findings indicate that NAc D2 neuron inhibition induces a negative hedonic value that an HFD choice test can unmask.

### Inhibition of NAc D2 neuron drives negative reinforcement

After establishing that optogenetic activation of NAc D2-expressing neurons induces place preference, we sought to determine whether inhibition of these neurons elicits a negative hedonic value using a negative reinforcement task. In this more sensitive paradigm, mice had to enter an ‘off’ zone to end optogenetic stimulation delivered in the “on” zone. Stimulation was applied at 20 Hz for the D2^ChR2^ and D2^tdTomato^ groups and continuously applied for the D2^ArchT^ group (**Fig 7a**). Despite the natural tendency of mice to explore, which led to most of their time being spent in the ‘on’ zone, the D2^ArchT^ group showed a significant shift in behavior, with increased time spent in the ‘off’ zone compared to the D2^tdTomato^ and D2^ChR2^ groups (**Figs 7b, 7c** [*one-way ANOVA; main factor: groups; F_(2, 33)_ = 4.27, p = 0.03]), indicating that sustained inhibition of NAc D2-expressing neurons generated a negative valence, prompting mice to avoid the area of stimulation.

**Figure 7.**
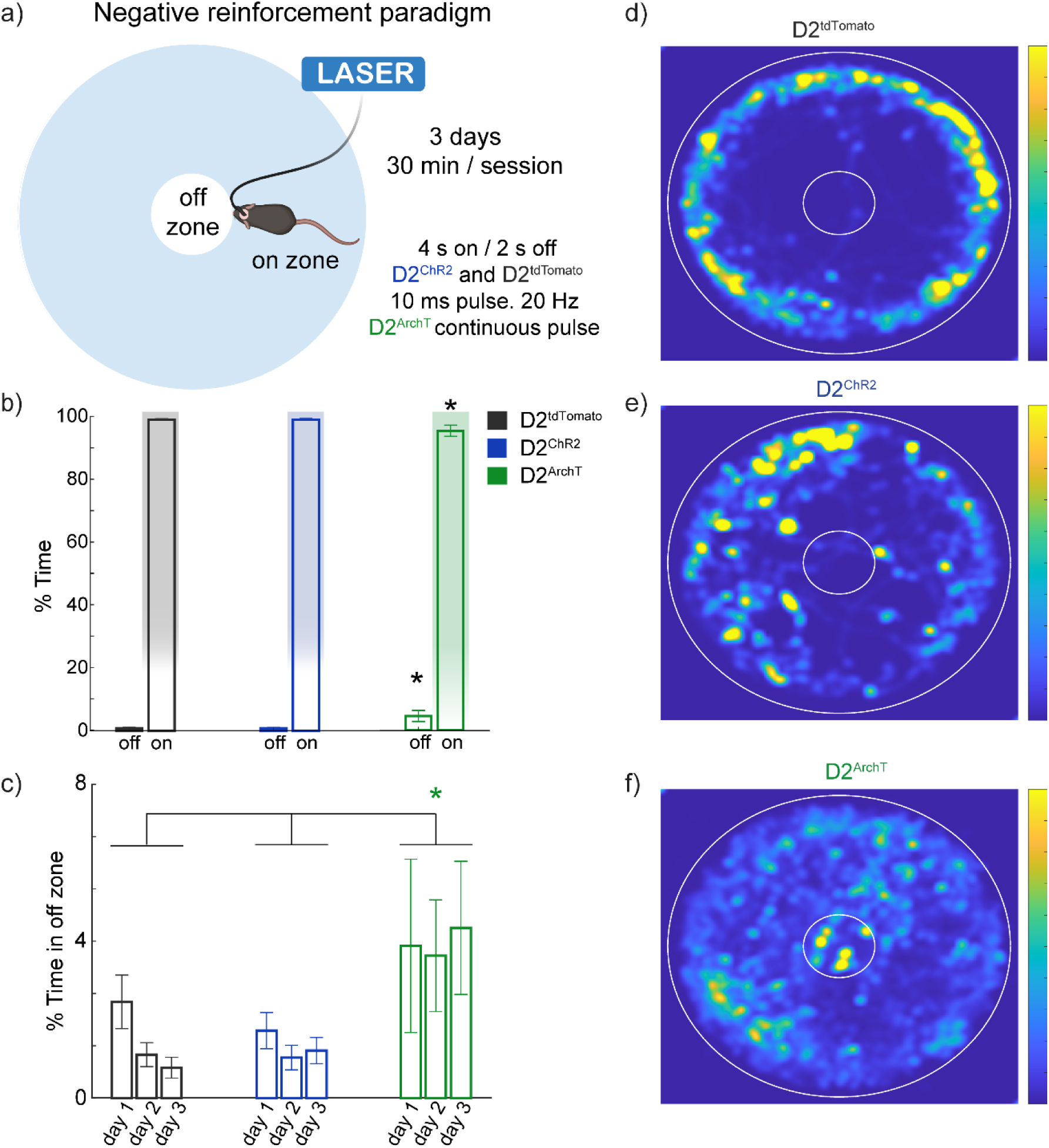
Silencing of NAc D2 neuron activity is hedonically negative and drives negative reinforcement. (**a**) Schematic illustration of the negative reinforcement paradigm. Mice were placed in a circular arena with distinct ‘on’ (optogenetic stimulation; 4 s on, 2 s off) and ‘off’ zones that turn off the stimulation. D2^ChR2^ and D2^tdTomato^ groups received pulsed stimulation in the ‘on’ zone, while D2ArchT received continuous stimulation. (**b**) Percentage of time spent in the ‘on’ and ‘off’ zones during the test for D2^tdTomato^ (gray), D2^ChR2^ (blue), and D2^ArchT^ (green) groups. (**c**) Time course of the percentage of time spent in the initial ‘off’ zone across the 3-day sessions. Mice expressing D2^ArchT^ showed increased time spent in the ‘off’ zone on day 3, indicative of negative reinforcement behavior. (**d-f**) Representative heatmaps showing the spatial distribution of time spent by mice in the D2^tdTomato^ (**d**), D2^ChR2^ (**e**), and D2^ArchT^ (**f**) groups during the post-conditioning test. * Significant difference compared to other groups. All significant differences had p < 0.05. Data are mean ± SEM. The statistical analyses were conducted with one-way ANOVA followed by Tukey-Kramer post hoc tests. D2^tdTomato^ n=6, D2^ChR2^ n=4, D2^ArchT^ n=4.

Representative heatmaps from the third session (**Figs 7d-f**) further illustrate this behavior, emphasizing the negative reinforcement effect of D2 neuron inhibition. These results show that inhibiting NAc D2 neurons elicits a negative hedonic value, with mice actively avoiding areas where stimulation occurs. This active avoidance behavior highlights the role of NAc D2-expressing neurons in driving negative reinforcement.

## Discussion

This study demonstrates that dopamine D2 receptor-expressing neurons in the NAc play a dual role in reward processing and ingestive behavior. Optogenetic activation of these neurons at higher frequencies disrupts the licking microstructure, decreasing sucrose preference while paradoxically increasing self-stimulation. This seemingly contradictory effect highlights the complex influence of D2 neuron activity, which can simultaneously enhance incentive motivation while disrupting specific oromotor behaviors. Interestingly, synchronizing stimulation with the animal’s natural licking rhythm minimized licking disruptions and promoted sucrose intake. Regarding spatial contextual reward, high-frequency stimulation (20 Hz) was sufficient to drive place preference (Fig 5f) and establish a long-term spatial preference memory (Fig 6d), confirming the rewarding properties of D2 neuron activation. Conversely, inhibiting these neurons led to a negative hedonic state (but not complete aversion), affecting food choices in specific contexts, particularly when a high-fat diet (HFD) was involved. These findings demonstrate the dual role of NAc D2-expressing neurons in regulating reward and consumption, highlighting the potential of entrained, self-paced stimulation that leverages the naturalistic licking theta rhythm to precisely activate these neurons.

### Activation of NAc D2 neurons reduces sucrose preference and increases self-stimulation

Optogenetic activation of NAc D2 neurons at 20 Hz disrupted sucrose preference by fragmenting the licking microstructure (**Fig. 1**). This finding is consistent with previous studies emphasizing the role of the NAc in orchestrating consummatory actions and processing palatability-related information.[8] In this regard, electrophysiological and calcium imaging studies in rats and mice have shown that a subset of NAc neurons encodes sucrose concentration and palatability, while other subsets exhibit oromotor responses entrained to the licking rhythm,[1,2,21,22] highlighting that the NAc is embedded in dynamic circuits that regulate hedonic processing and goal-directed behaviors.[7] While D1 neurons have traditionally been associated with feeding,[8,9] our findings indicate that experimentally imposed 20 Hz activation of D2 neurons particularly disrupts ingestive behavior.

### Frequency-dependent disruption of licking behavior by NAc D2-expressing neurons

Prado et al. (2016) showed for the first time that optogenetic manipulations of glutamatergic inputs to the NAc are rewarding but also stopped licking behavior in a frequency-dependent manner, reducing lick duration and frequency, resembling our observation of D2 neurons activation shortened lick bout durations and reduced bout sizes (**Fig 2**).[14] Interestingly, these disruptions were frequency-dependent, with higher frequencies (15 and 20 Hz) producing more pronounced effects, supporting the notion that NAc D2-expressing neurons would be, at least, one target of glutamatergic inputs to fine-tune the intensity and timing of consummatory actions.[14]

In a detailed frequency test, we found that higher optogenetic stimulation frequencies (15 and 20 Hz) produced a significant suppression of licking behavior compared to lower frequencies. These results support that D2-expressing neurons act as modulators of consummatory actions depending on the stimulation pattern, as highlighted by Soares-Cunha et al. (2019). In their study, brief stimulation was rewarding, but prolonged stimulation became aversive and induced more pronounced behavioral impairments.[5] Our study suggests that D2-expressing neurons play a dual role in regulating consummatory behavior. While they enhance the incentive motivation to self-stimulate, they simultaneously impair the execution of licking, as shown by increased self-stimulation despite the paradoxical impairment of licking (**Fig 4**). These results also indicate that NAc D2-expressing neurons can modulate ingestive behavior through frequency-dependent mechanisms, aligning with the concept of the NAc as a central hub integrating reward and feeding behaviors, where the activity of D2-expressing neurons can either promote or hinder consumption depending on the specific stimulation pattern.[8,14,15]

Our data can also reconcile some differences in literature. For example, O’Connor et al. (2015) found that 20 Hz stimulation of D1 neurons abruptly impaired licking, [9] whereas Sandoval et al. (2023) did not observe licking impairment using a lick-paired self-stimulation protocol of D1 neurons, like the one used in this study. Surprisingly, like us, they also observed a slight licking impairment using lick-paired self-stimulation of D2-expressing neurons, [11] suggesting a unique property of self-rhythmic licking to stimulate the NAc. These results also support the notion that rhythmic licking acts as an internal clock to synchronize brain activity across different brain regions.[23] While our findings show that D2-expressing neurons impair rhythmic licking, particularly at high frequencies, the precise circuit mechanisms underlying this disruption remain unclear. This is because previous research has shown that lesioning D1, but not D2, neurons impairs licking microstructure,[11] and rabies tracing studies indicate that, unlike D1 neurons, D2 neurons do not directly connect to taste buds or the CPG for licking.[11] One possibility is that D2-expressing neurons disrupt licking behavior indirectly by modulating D1 neuron activity, potentially through lateral inhibition.[24,25] This hypothesis warrants further investigation, which could employ cell-type-specific manipulations and circuit-tracing techniques to delineate the precise pathways and mechanisms involved.

### Self-paced stimulation enhances task engagement and reduces licking disruption

One of the key insights from the present study is the better preservation of licking behavior and enhanced task engagement when stimulation was synchronized with natural licking rhythms (**Figs 3,4**), highlighting the flexible nature of reward-seeking behaviors governed by the NAc. This finding aligns with previous research showing that mice will self-stimulate glutamatergic inputs to the NAc and that these glutamatergic inputs are suppressed during feeding.[14,26] This suggests that NAc circuits dynamically adapt to regulate both consumption and the pursuit of rewards.[13,14,26] Similarly, Cole et al. (2018) reported that allowing animals to self-stimulate D2 neurons by licking a spout, mirroring their natural behavior, led to increased engagement and fewer disruptions compared to externally imposed stimulation triggered by entering a specific spatial location.[17] These findings resonate with the concept that D2 neuron activity can modulate reward by integrating motor information into reward-seeking,[27] promoting goal-directed behaviors while maintaining hedonic value.[1] This supports the view that stimulation parameters, particularly self-paced stimulation, are crucial in determining the behavioral outcomes of D2 neuron activation.[4]

### High-frequency activation of NAc D2 neurons drives place preference and enhances exploratory behavior

In addition to modulating licking behaviors, our study revealed that high-frequency activation of NAc D2 neurons (20 Hz) also induced robust place preference and significantly enhanced exploratory behavior, indicating a strong, rewarding effect (**Fig 5**). While place preference has traditionally been associated with D1 neurons,[19] our results extend these findings to D2-expressing neurons, indicating that these neurons play a more significant role in reward-seeking behaviors than previously assumed. Our data also supports the observation of Cole et al. (2018) using a lick-spout self-stimulation task. They found that mice self-stimulated both D1 and D2 neurons in NAc, although D1 excitation was more rewarding than D2 neurons. They also suggest that the excitation of D2 neurons in NAc supports self-stimulation under some tasks (using a lick-spout) but fails under other more complex and demanding tasks.[17] Oginsky et al. (2016) showed that NAc D2-expressing neurons are involved in both place preference and exploratory behaviors, particularly in contexts involving palatable food, supporting our findings of enhanced exploratory drive during 20 Hz stimulation.[28]

The enhanced exploratory behavior observed in the stimulated context suggests that D2-expressing neurons may contribute to a broader behavioral repertoire beyond reward.[15] This indicates a potential role for D2-expressing neurons in modulating spatial exploration and context-dependent behaviors, contributing to the animal’s overall motivational state during task engagement.[25]

Previous studies using pharmacological, dopaminergic projections, or optogenetic perturbations have reported the existence of hotspots in which dedicated subregions of the NAc play different roles in reward and aversion.[29–31]. An elegant study by Chen et al. (2023) recently demonstrated that optogenetic activation of neurotensin (Nts+) neurons (80% D1 and 20% D2) in the lateral NAc shell at 20 Hz elicits reward, while activation of cocaine- and amphetamine-regulated transcript prepropeptide (Cartpt+) neurons (comprising an equal subset of both D1 and D2 MSNs) in the medial NAc shell is aversive. This finding appears to contradict our results and others,[5,17,25,32] which indicate that 20 Hz activation of D2 neurons in the medial NAc is rewarding. Likewise, our findings further challenge the traditional dichotomy that assigns reward facilitation to D1-expressing neurons and aversion mediation to D2-expressing neurons.[16,18,33] Instead, NAc D2 neurons could participate in positive and negative valence depending on stimulation parameters and behavioral context. This apparent contradiction may be reconciled by considering the multifaceted nature of NAc function. As highlighted here, the role of NAc neurons appears to be not solely determined by cell type or anatomical location but is also critically influenced by stimulation parameters and behavioral context.

Our results are consistent with studies that indicate the importance of context and stimulation parameters in shaping the behavioral outcomes of NAc D2 neuron activation and inhibition.[3,5,14,17]

### Inhibition of NAc D2 neurons decreases palatable food intake and drives context-dependent negative reinforcement

Interestingly, while optogenetic activation produced place preference, inhibition of D2 neurons also revealed a role in processing hedonic value (**Fig 6**). Specifically, inhibition led to a preference for HFD consumption on the non-stimulated side of a conditioned chamber, suggesting that D2 neuron suppression induces a negative hedonic value that palatable food can partially mask. Yang et al. (2020) found that inhibition of D2 neurons unmasked negative hedonic value in specific contexts, mirroring our findings that D2 inhibition during HFD reveals a negative hedonic signal that palatable food can override.[3] Cole et al. (2018) reported that optogenetic activation of D2 neurons leads to a motivational state where both reward-seeking and sometimes negative avoidance responses coexist, suggesting that D2-expressing neurons exhibit a context-dependent ambivalence response.[17] These findings are consistent with earlier studies demonstrating that D2 neurons contribute to positive and negative valence processing depending on the context and task structure.[4,5] This result emphasizes the context-dependent nature of D2 neuron function, where bulk inhibition generates a negative hedonic state, further supporting the idea that D2-expressing neurons can induce a negative reinforcement (**Fig 7**). Our findings extend the current understanding of NAc D2-expressing neurons, showing their critical role in balancing reward and consummatory behavior across diverse contexts of motivation and motor control. This duality highlights their complex contribution to reward circuits, aligning with the broader literature on their dual functions.[15,17,34]

### Limitation of the study

We acknowledge that the attenuated efficacy of D2 inhibition observed in this study may be attributed, in part, to the low spontaneous firing rate of NAc D2 neurons (<1 Hz; [35]), suggesting a ‘floor effect’ whereby further suppression has minimal impact.[36–38] Nevertheless, we observed that D2 inhibition induced a clear negative hedonic value, demonstrating the efficacy of this brain perturbation. However, ChR2-mediated activation, particularly at high frequencies, may induce supraphysiological firing patterns that exceed the low endogenous activity of D2 neurons, thereby amplifying behavioral effects.[14] Our observation that stimulation phase-locked to self-paced licking has little impact on licking, whereas high-frequency, experimenter-imposed stimulation suppresses it, highlights a need for further studies to uncover the neuronal correlates underlying this interesting phenomenon. While this study did not record the natural firing rate of D2 neurons during licking, previous work in our lab and others.[2,21] Cho et al., (2007), Gutierrez et al., (2010), and Villavicencio et al., (2018) revealed a subset of rat NAc neurons that fire in coherence with rhythmic licking behavior.[2,23,39] This suggests that self-paced stimulation may further entrain and amplify lick-coherent activity within this population of NAc neurons, thereby mitigating the impact on licking behavior. In contrast, high-frequency stimulation (e.g., 20 Hz) imposes firing patterns that exceed physiological activity,[35,40,41] likely disrupting rhythmic coordination and licking microstructure.

Although our findings provide new insights into the role of NAc D2-expressing neurons in reward and consummatory behaviors, it is important to acknowledge some limitations. Firstly, while this study focused on D2-expressing neurons, future research should investigate the role of D1-expressing neurons with similar paradigms to understand better the network dynamics governing these behaviors. Future work should explore how D1 and D2 neurons interact to regulate both homeostatic and hedonic feeding, building upon work by Walle et al. (2024), using simultaneous manipulations of D1 and D2 expressing neurons that highlighted distinct but complementary roles for these neurons in energy balance and reward-seeking.[15] Secondly, while we observed significant frequency-dependent effects, the precise circuits and downstream targets mediating these effects require further investigation. Thirdly, our experiments did not include a condition where water was the only available stimulus, which limits our ability to generalize the observed suppression of consummatory behavior to non-palatable rewards. Future research should address whether D2-ChR2 stimulation disrupts consummatory actions in general, specifically targets hedonic reward processing, or affects a combination of both. It is important to recognize that our use of optogenetics to modulate NAc D2-expressing neurons (or any other cell type) while providing valuable insights may not fully capture the ‘physiological’ role of endogenous neural activity and naturalistic reward behaviors.[41] This study found that behavioral responses are highly dependent on optogenetic stimulation parameters. Therefore, chemogenetic manipulations are also expected to produce varying behavioral effects as different stimulation approaches engage neuronal circuits in distinct ways.[15] The observation that different stimulation parameters of D2-expressing neurons result in different behavioral outputs aligns with the concept of optoception. We recently reported the optoception phenomenon, suggesting that mice could perceive each stimulation pattern slightly differently, providing a basis for mice’s ability to identify and discriminate between various optogenetic stimulation frequencies.[41] Finally, because cholinergic interneurons in the NAc also express D2 receptors, future studies employing intersectional genetic techniques are needed to differentiate the contribution of these interneurons from those of D2 MSNs. This will provide a more precise understanding of the specific roles of these distinct cell populations in reward and consumption.

## Conclusion

In summary, our findings demonstrate the dual role of NAc D2-expressing neurons in modulating reward and consummatory behavior, where activation can simultaneously enhance incentive motivation to self-stimulate and disrupt licking behavior. These results highlight the importance of stimulation parameters, frequency, timing, and agency of actions in shaping the behavioral outcomes of D2 neuron activity. Inhibition of these neurons induces a negative hedonic state that is (un)masked in specific contexts, such as during HFD consumption. Thus, NAc D2-expressing neurons act as critical modulators, balancing reward and licking disruption through rhythmic optogenetic stimulation, providing new insights into their role in reward circuits. These findings deepen our understanding of the NAc role in reward circuits and may provide potential avenues for exploring therapeutic interventions for disorders related to reward processing, such as addiction[42] or obesity.[43]

## Data Availability Statement

All relevant data are included in the paper and its supporting information files. Additional data is available on request from the corresponding author.

## Acknowledgments

All behavioral task schemes were created using BioRender. We thank Elvi Gil Lievana for her help with histology, Fabiola Olvera Hernandez for her valuable assistance with animal care, and Jorge Luis-Islas for his collaboration in the early stages of this project. We also appreciate the comments from Luis Tellez on a previous version of this manuscript.

## Supporting Information

**S1 Table. Microstructure analysis of licking for two-bottle sucrose test, Fig 1.**

**S2 Table. Microstructure analysis of licking for sucrose freely-licking with optogenetic frequency scan test for Fig 2.**

**S3 Table. Microstructure analysis of licking for sucrose brief access taste test with frequency scan test for Fig 3.**

**S4 Table. Microstructure analysis of licking for sucrose brief access taste test delay scan test for Fig 4.**

**S5 Table. Statistical analysis**

