## Supplementary material for "Nucleus accumbens D2-expressing neurons: Balancing reward and licking disruption through rhythmic optogenetic stimulation": S1 Table

**Supplementary Table 1. Analysis of licking microstructure for the two-bottle sucrose test (Fig 1).**

| | | $D2^{tdTomato}$ Water | $D2^{tdTomato}$ Sucrose | $D2^{ChR2}$ Water | $D2^{ChR2}$ Sucrose | $D2^{ArchT}$ Water | $D2^{ArchT}$ Sucrose |
|---|---|---|---|---|---|---|---|
| Condition 1 | Base line (no-laser) | | | | | | |
| | Preference (%) | 21.69 ± 4.35 | 78.31 ± 4.35 | 11.96 ± 1.68 | 88.04 ± 1.68 | 14.53 ± 3.49 | 85.48 ± 3.49 |
| | Inter lick interval (ms) | 135.08 ± 3.35 | 130.32 ± 2.58 | 146.00 ± 5.99 | 134.43 ± 4.41 | 133.64 ± 2.90 | 128.94 ± 1.85 |
| | Lick rate (Hz) | 10.04 ± 0.47 | 9.64 ± 0.38 | 9.75 ± 0.39 | 9.50 ± 0.13 | 10.15 ± 0.52 | 9.40 ± 0.27 |
| | Bout size (Licks) | 14.10 ± 3.53 | 23.18 ± 5.33 | 9.89 ± 1.61 | 19.41 ± 2.03 | 12.96 ± 2.46 | 28.38 ± 4.57 |
| | Bout duration (s) | 1.71 ± 0.44 | 2.88 ± 0.69 | 1.23 ± 0.21 | 2.40 ± 0.26 | 1.60 ± 0.33 | 3.55 ± 0.61 |
| | Inter bout interval (s) | 82.49 ± 11.96 | 39.96 ± 3.88 | 101.50 ± 25.40 | 22.18 ± 2.45 | 101.34 ± 24.55 | 29.76 ± 2.88 |
| | Number of bouts | 14.31 ± 2.09 | 35.54 ± 3.52 | 17.86 ± 2.54 | 70.14 ± 7.58 | 14.94 ± 2.77 | 48.82 ± 6.89 |
| | Total licks | 182.92 ± 39.79 | 714.00 ± 119.42 | 152.41 ± 24.99 | 1140.09 ± 94.84 | 154.24 ± 34.42 | 1198.82 ± 154.07 |
| | Total licks sum | 896.92 ± 132.45 | | 1292.50 ± 101.87 | | 1353.06 ± 150.57 | |
| Condition 2 | Laser sucrose | | | | | | |
| | Preference (%) | 16.32 ± 3.70 | 83.68 ± 3.70 | **#*40.43 ± 4.28** | **#*59.57 ± 4.28** | 20.84 ± 4.48 | 79.16 ± 4.48 |
| | Inter lick interval (ms) | 137.22 ± 3.40 | 130.23 ± 1.83 | 136.46 ± 2.75 | **#*160.19 ± 4.25** | 137.23 ± 4.54 | 128.70 ± 3.65 |
| | Lick rate (Hz) | 9.59 ± 0.63 | 9.08 ± 0.25 | 9.17 ± 0.29 | ***9.86 ± 0.23** | **#8.86 ± 0.36** | 9.57 ± 0.26 |
| | Bout size (Licks) | 20.14 ± 4.61 | 32.34 ± 3.65 | **#22.68 ± 2.05** | **#*5.66 ± 0.57** | 29.22 ± 7.40 | 25.74 ± 4.02 |
| | Bout duration (s) | 2.59 ± 0.61 | 4.09 ± 0.47 | **#2.93 ± 0.29** | **#*0.72 ± 0.08** | 3.65 ± 0.95 | 3.20 ± 0.53 |
| | Inter bout interval (s) | 172.61 ± 85.84 | 45.11 ± 5.54 | 52.88 ± 5.41 | **#*14.78 ± 1.66** | 64.72 ± 24.94 | ***28.36 ± 3.43** |
| | Number of bouts | 9.00 ± 1.68 | 32.78 ± 4.03 | ***18.25 ± 1.66** | **#*121.21 ± 14.65** | 13.91 ± 2.80 | ***64.04 ± 10.03** |
| | Total licks | 172.61 ± 44.05 | 939.39 ± 123.44 | **#*414.42 ± 45.60** | **#659.25 ± 91.50** | 243.65 ± 45.51 | 1298.61 ± 166.53 |
| | Total licks sum | 1112.00 ± 122.96 | | 1073.67 ± 103.77 | | 1542.26 ± 159.17 | |
| Condition 3 | Laser water | | | | | | |
| | Preference (%) | 20.63 ± 5.18 | 79.37 ± 5.18 | 13.34 ± 3.34 | 86.66 ± 3.34 | **9.57 ± 1.80** | **90.43 ± 1.80** |
| | Inter lick interval (ms) | 145.94 ± 12.51 | 132.34 ± 2.61 | **#166.96 ± 7.05** | 134.50 ± 1.89 | 139.97 ± 10.09 | 128.62 ± 2.22 |
| | Lick rate (Hz) | 10.50 ± 0.72 | 9.09 ± 0.35 | 9.38 ± 0.38 | **#8.71 ± 0.21** | 9.93 ± 0.57 | 9.71 ± 0.28 |
| | Bout size (Licks) | 17.44 ± 4.60 | 31.50 ± 5.04 | **#5.15 ± 1.49** | **#30.51 ± 3.55** | 14.14 ± 6.83 | 22.69 ± 3.98 |
| | Bout duration (s) | 2.18 ± 0.59 | 4.02 ± 0.69 | **0.65 ± 0.23** | **#3.93 ± 0.44** | 1.70 ± 0.87 | 2.76 ± 0.50 |
| | Inter bout interval (s) | 168.50 ± 42.56 | 61.76 ± 5.59 | **#*24.90 ± 2.91** | ***28.28 ± 4.61** | ***78.05 ± 12.98** | **#*20.22 ± 2.93** |
| | Number of bouts | 9.42 ± 1.53 | 23.92 ± 2.80 | **#*58.38 ± 14.21** | ***61.69 ± 8.02** | 17.82 ± 4.70 | ***78.82 ± 17.14** |
| | Total licks | 161.00 ± 41.41 | 647.25 ± 93.14 | 245.38 ± 66.20 | **#*1588.12 ± 137.07** | 132.73 ± 24.25 | ***1475.27 ± 223.91** |
| | Total licks sum | 808.25 ± 89.56 | | **#*1833.50 ± 142.29** | | ***1608.00 ± 231.36** | |

Bold font indicates statistically significant differences ($p<0.05$):

*****Significant difference compared to the $D2^{tdTomato}$ (control group) in the same condition.

#Significant difference compared to Baseline (condition 1) within the same group.
