## Supplementary material for "Nucleus accumbens D2-expressing neurons: Balancing reward and licking disruption through rhythmic optogenetic stimulation": S2 Table

**Supplementary Table 2. Analysis of licking microstructure for sucrose freely licking with optogenetic frequency scan test (Fig 2).**

| Stimualtion Frequencies | | $D2^{tdTomato}$ | $D2^{ChR2}$ | $D2^{ArchT}$ |
|---|---|---|---|---|
| No-laser | Inter lick interval (ms) | 159.75 ± 7.40 | 168.31 ± 5.45 | 162.87 ± 5.26 |
| | Lick rate (Hz) | 8.08 ± 0.23 | 8.21 ± 0.40 | 7.88 ± 0.15 |
| | Bout size (Licks) | 5.41 ± 0.37 | 4.73 ± 0.15 | 5.31 ± 0.18 |
| | Bout duration (s) | 0.68 ± 0.03 | 0.61 ± 0.03 | 0.69 ± 0.01 |
| | Inter bout interval (s) | 159.01 ± 16.82 | 197.99 ± 19.95 | 152.97 ± 13.27 |
| | Number of bouts | 7.29 ± 0.81 | 6.52 ± 0.47 | 8.62 ± 1.16 |
| | Total licks | 40.04 ± 3.46 | 32.37 ± 2.59 | 49.42 ± 8.00 |
| 5 Hz | Inter lick interval (ms) | 176.21 ± 16.76 | **#202.76 ± 8.79** | 165.96 ± 5.21 |
| | Lick rate (Hz) | 7.56 ± 0.26 | 7.74 ± 0.28 | 7.75 ± 0.26 |
| | Bout size (Licks) | 4.68 ± 0.32 | **#3.69 ± 0.27** | 5.15 ± 0.19 |
| | Bout duration (s) | 0.61 ± 0.03 | **#0.50 ± 0.03** | ***0.67 ± 0.01** |
| | Inter bout interval (s) | 153.97 ± 28.50 | 215.33 ± 38.07 | 169.83 ± 21.92 |
| | Number of bouts | 7.44 ± 0.89 | 6.56 ± 0.47 | 8.36 ± 1.15 |
| | Total licks | 36.39 ± 5.61 | 25.61 ± 1.92 | 46.08 ± 7.59 |
| 8 Hz | Inter lick interval (ms) | 168.78 ± 9.86 | **#*234.59 ± 11.70** | 171.09 ± 6.46 |
| | Lick rate (Hz) | 8.09 ± 0.34 | 7.27 ± 0.49 | 7.72 ± 0.21 |
| | Bout size (Licks) | 4.50 ± 0.46 | **#*2.81 ± 0.21** | 4.69 ± 0.24 |
| | Bout duration (s) | 0.57 ± 0.05 | **#*0.39 ± 0.04** | **#0.61 ± 0.02** |
| | Inter bout interval (s) | 126.58 ± 19.68 | 206.92 ± 24.93 | 151.09 ± 19.49 |
| | Number of bouts | 7.55 ± 0.78 | 5.82 ± 0.51 | 8.23 ± 1.20 |
| | Total licks | 36.59 ± 6.06 | **#*17.73 ± 2.04** | 42.43 ± 7.51 |
| 15 Hz | Inter lick interval (ms) | 156.65 ± 7.71 | **#*221.38 ± 12.27** | 171.81 ± 5.07 |
| | Lick rate (Hz) | 8.28 ± 0.43 | 7.50 ± 0.37 | 7.55 ± 0.22 |
| | Bout size (Licks) | 4.93 ± 0.19 | **#*2.70 ± 0.52** | 4.79 ± 0.22 |
| | Bout duration (s) | 0.60 ± 0.01 | **#0.33 ± 0.09** | 0.64 ± 0.02 |
| | Inter bout interval (s) | 126.25 ± 17.34 | 166.18 ± 31.86 | 180.49 ± 41.11 |
| | Number of bouts | 7.69 ± 1.03 | ***5.12 ± 0.58** | 8.30 ± 1.16 |
| | Total licks | 39.59 ± 4.37 | **#*15.56 ± 3.48** | 43.77 ± 7.66 |
| 21 Hz | Inter lick interval (ms) | 168.95 ± 9.95 | **#*214.47 ± 10.03** | 166.83 ± 5.49 |
| | Lick rate (Hz) | 7.96 ± 0.39 | 8.33 ± 0.63 | 8.13 ± 0.22 |
| | Bout size (Licks) | 4.76 ± 0.07 | **#*2.41 ± 0.55** | **#4.52 ± 0.20** |
| | Bout duration (s) | 0.62 ± 0.03 | **#*0.28 ± 0.09** | **#0.57 ± 0.02** |
| | Inter bout interval (s) | 146.22 ± 20.15 | 209.07 ± 38.68 | 167.95 ± 20.98 |
| | Number of bouts | 7.30 ± 1.15 | 4.92 ± 0.84 | 8.37 ± 1.09 |
| | Total licks | 35.47 ± 6.25 | **#*13.28 ± 3.47** | 41.75 ± 6.92 |

Bold font indicates statistically significant differences ($p<0.05$):

*****Significant difference compared to the $D2^{tdTomato}$ (control group) at the same frequency.

#Significant difference compared to the Control (no laser) frequency within the same group.
