## Supplementary material for "Nucleus accumbens D2-expressing neurons: Balancing reward and licking disruption through rhythmic optogenetic stimulation": S3 Table

**Supplementary Table 3. Analysis of licking microstructure for suc**
**frequency scan test (Fig 3).**

| Stimulation pattern | | | $D2^{tdTomato}$ | $D2^{ChR2}$ |
|---|---|---|---|---|
| Stimulation pattern | No-laser | Inter lick interval (ms) | 171.05 ± 7.13 | 168.30 ± 8.88 |
| | | Lick rate (Hz) | 4.02 ± 0.26 | 4.10 ± 0.47 |
| | | Bout size (licks) | 15.73 ± 1.04 | 16.00 ± 1.97 |
| | | Bout duration (s) | 2.40 ± 0.12 | 2.33 ± 0.25 |
| | | Inter bout interval (s) | 129.94 ± 11.21 | ***83.82 ± 9.69** |
| | | Number of bouts | 8.00 ± 1.02 | ***16.10 ± 2.14** |
| | | Total licks | 124.77 ± 22.91 | ***214.71 ± 23.36** |
| | Lick paired | Inter lick interval (ms) | 174.78 ± 9.90 | 190.00 ± 8.97 |
| | | Lick rate (Hz) | 4.13 ± 0.38 | 3.54 ± 0.32 |
| | | Bout size (licks) | 16.04 ± 1.56 | 13.53 ± 1.43 |
| | | Bout duration (s) | 2.37 ± 0.17 | 2.24 ± 0.22 |
| | | Inter bout interval (s) | 152.61 ± 18.14 | ***79.15 ± 9.33** |
| | | Number of bouts | 8.68 ± 1.29 | ***16.14 ± 1.89** |
| | | Total licks | 127.09 ± 24.92 | 187.14 ± 18.81 |
| | 8 Hz | Inter lick interval (ms) | 172.28 ± 8.50 | 187.64 ± 9.91 |
| | | Lick rate (Hz) | 4.00 ± 0.37 | **@#*2.54 ± 0.29** |
| | | Bout size (licks) | 15.13 ± 1.62 | **#*9.89 ± 1.18** |
| | | Bout duration (s) | 2.24 ± 0.20 | **@#*1.55 ± 0.17** |
| | | Inter bout interval (s) | 129.70 ± 14.18 | ***68.95 ± 8.01** |
| | | Number of bouts | 8.82 ± 1.22 | ***18.38 ± 2.15** |
| | | Total licks | 119.91 ± 21.86 | **#150.19 ± 15.43** |
| | 20 Hz | Inter lick interval (ms) | 171.02 ± 8.56 | 180.40 ± 9.14 |
| | | Lick rate (Hz) | 4.16 ± 0.36 | **#*2.81 ± 0.40** |
| | | Bout size (licks) | 16.44 ± 1.46 | **@#*8.64 ± 1.66** |
| | | Bout duration (s) | 2.46 ± 0.16 | **@#*1.30 ± 0.28** |
| | | Inter bout interval (s) | 142.75 ± 13.16 | ***87.00 ± 9.26** |
| | | Number of bouts | 7.36 ± 0.88 | ***14.57 ± 1.58** |
| | | Total licks | 119.64 ± 22.17 | **!@#97.14 ± 14.57** |

Bold font indicates statistically significant differences ($p < 0.05$):

***** Significant difference compared to the $D2^{tdTomato}$ (control group) at the sar

\# Significant difference compared to the Control (no laser) frequency withir

@ Significant difference between the Lick-paired condition and the 8 Hz or within the same group

**!** Significant difference between the 8 Hz and 20 Hz conditions within the s rose BATT with

| D2$^{ArchT}$ |
|---|
| ***150.41 ± 4.78** |
| ***5.04 ± 0.32** |
| ***20.00 ± 1.28** |
| 2.82 ± 0.17 |
| 178.04 ± 33.50 |
| 6.80 ± 0.62 |
| 122.36 ± 7.36 |
| ***148.58 ± 3.94** |
| 4.55 ± 0.29 |
| 18.07 ± 1.14 |
| 2.49 ± 0.15 |
| 150.76 ± 16.10 |
| 7.16 ± 0.52 |
| 121.16 ± 7.88 |
| ***143.06 ± 3.34** |
| ***5.29 ± 0.31** |
| ***20.88 ± 1.26** |
| ***2.81 ± 0.16** |
| 159.22 ± 15.26 |
| 6.48 ± 0.58 |
| 125.20 ± 8.88 |
| ***144.91 ± 4.74** |
| 5.13 ± 0.40 |
| 18.80 ± 1.64 |
| 2.51 ± 0.19 |
| 175.73 ± 30.39 |
| 6.72 ± 0.56 |
| 121.04 ± 10.41 |

ne frequency
the same group
20 Hz conditions

ame group
