## Supplementary material for "Nucleus accumbens D2-expressing neurons: Balancing reward and licking disruption through rhythmic optogenetic stimulation": S4 Table

**Supplementary Table 4. Analysis of licking microstructure analysis of licking for the sucrose BATT with varying optogenetic stimulation delay (Fig 4).**

| Stimulation pattern | | | $D2^{tdTomato}$ | $D2^{ChR2}$ | $D2^{ArchT}$ |
|---|---|---|---|---|---|
| | No-laser | Inter lick interval (ms) | 150.95 ± 5.76 | ***179.48 ± 8.71** | 138.06 ± 4.97 |
| | | Lick rate (Hz) | 5.04 ± 0.36 | ***3.48 ± 0.57** | 5.26 ± 0.45 |
| | | Bout size (licks) | 19.89 ± 1.50 | ***13.85 ± 2.30** | 20.75 ± 1.80 |
| | | Bout duration (s) | 2.80 ± 0.22 | 2.18 ± 0.36 | 2.74 ± 0.28 |
| | | Inter bout interval (s) | 191.64 ± 21.62 | ***84.71 ± 11.81** | 200.87 ± 38.04 |
| | | Number of bouts | 5.60 ± 0.72 | ***15.70 ± 2.97** | 5.50 ± 1.09 |
| | | Total licks | 110.07 ± 19.52 | 181.00 ± 36.21 | 111.12 ± 21.37 |
| | 0 ms delay (lick paired) | Inter lick interval (ms) | 156.21 ± 6.95 | ***206.89 ± 13.27** | 134.84 ± 5.75 |
| | | Lick rate (Hz) | 4.76 ± 0.44 | ***2.68 ± 0.46** | 5.23 ± 0.64 |
| | | Bout size (licks) | 18.74 ± 1.76 | ***10.53 ± 1.85** | 20.78 ± 2.55 |
| | | Bout duration (s) | 2.65 ± 0.19 | ***1.81 ± 0.27** | 2.58 ± 0.26 |
| | | Inter bout interval (s) | 206.87 ± 27.54 | ***79.25 ± 12.82** | 174.32 ± 20.18 |
| | | Number of bouts | 6.20 ± 0.94 | ***18.20 ± 3.54** | 4.88 ± 0.74 |
| | | Total licks | 110.73 ± 19.11 | 159.10 ± 32.62 | 96.75 ± 14.28 |
| | 50 ms delay | Inter lick interval (ms) | 151.28 ± 8.05 | ***190.58 ± 8.62** | 149.19 ± 5.76 |
| | | Lick rate (Hz) | 4.57 ± 0.43 | ***2.88 ± 0.45** | 5.12 ± 0.75 |
| | | Bout size (licks) | 18.16 ± 1.74 | ***11.36 ± 1.83** | 20.22 ± 3.03 |
| | | Bout duration (s) | 2.48 ± 0.21 | 1.87 ± 0.29 | 2.76 ± 0.34 |
| | | Inter bout interval (s) | 209.14 ± 29.89 | ***82.81 ± 12.84** | 171.57 ± 35.85 |
| | | Number of bouts | 6.27 ± 0.75 | ***18.00 ± 3.81** | 5.88 ± 0.79 |
| | | Total licks | 105.87 ± 16.64 | 167.70 ± 34.42 | 114.12 ± 19.19 |
| | 100ms delay | Inter lick interval (ms) | 147.26 ± 5.72 | ***188.82 ± 7.84** | 139.84 ± 7.25 |
| | | Lick rate (Hz) | 4.82 ± 0.44 | ***3.32 ± 0.57** | 6.39 ± 0.71 |
| | | Bout size (licks) | 18.87 ± 1.83 | 13.26 ± 2.30 | 25.14 ± 2.84 |
| | | Bout duration (s) | 2.53 ± 0.23 | 2.21 ± 0.38 | 3.25 ± 0.28 |
| | | Inter bout interval (s) | 212.17 ± 21.39 | ***84.43 ± 13.25** | 148.41 ± 20.66 |
| | | Number of bouts | 5.47 ± 0.72 | ***17.00 ± 3.47** | 4.88 ± 0.52 |
| | | Total licks | 97.00 ± 14.35 | ***178.50 ± 32.34** | 119.88 ± 17.51 |

Bold indicates statistically significant differences ($p < 0.05$):

***** Significant difference compared to the $D2^{tdTomato}$ (control group) in the same condition (delay time).
