## Supplementary material for "Nucleus accumbens D2-expressing neurons: Balancing reward and licking disruption through rhythmic optogenetic stimulation": S5 Table

### Fig 6 a

| | Group1 | Group2 | Condition | T-value | df |
|---|---|---|---|---|---|
| T-test | Control 20Hz | D2ArchT Continuous | Session 1 | -0.82370182 | 20 |
| T-test | Control 20Hz | D2ArchT Continuous | Session 2 | -0.82370182 | 20 |
| T-test | D2ArchT | - | Across Sessions | -3.06889597 | 11 |

### Fig 6 b

| | Group1 | Group2 | Condition | F-value | df1 |
|---|---|---|---|---|---|
| ANOVA | All groups | - | - | 1.543190053 | 2 |

### Fig 6d

| | Group1 | Group2 | Condition | F_value | df1 |
|---|---|---|---|---|---|
| ANOVA | All groups | | Pre-test | 13.17456071 | 2 |
| Tukey | D2tdTomato | D2ChR2 | | | |
| Tukey | D2tdTomato | D2ArchT | | | |
| Tukey | D2ChR2 | D2ArchT | | | |
| ANOVA | All groups | | Acq1 | 6.052216787 | 2 |
| Tukey | D2tdTomato | D2ChR2 | | | |
| Tukey | D2tdTomato | D2ArchT | | | |
| Tukey | D2ChR2 | D2ArchT | | | |
| ANOVA | All groups | | Acq2 | 17.60887331 | 2 |
| Tukey | D2tdTomato | D2ChR2 | | | |
| Tukey | D2tdTomato | D2ArchT | | | |
| Tukey | D2ChR2 | D2ArchT | | | |
| ANOVA | All groups | | Acq3 | 8.194371097 | 2 |
| Tukey | D2tdTomato | D2ChR2 | | | |
| Tukey | D2tdTomato | D2ArchT | | | |
| Tukey | D2ChR2 | D2ArchT | | | |
| ANOVA | All groups | | test | 11.23717169 | 2 |
| Tukey | D2tdTomato | D2ChR2 | | | |
| Tukey | D2tdTomato | D2ArchT | | | |
| Tukey | D2ChR2 | D2ArchT | | | |
| ANOVA | All sessions | D2tdTomato | | 0.870556542 | 4 |
| Tukey | Pre-test | Acq1 | | - | - |
| Tukey | Pre-test | Acq2 | | - | - |
| Tukey | Pre-test | Acq3 | | - | - |
| Tukey | Pre-test | test | | - | - |
| Tukey | Acq1 | Acq2 | | - | - |
| Tukey | Acq1 | Acq3 | | - | - |
| Tukey | Acq1 | test | | - | - |
| Tukey | Acq2 | Acq3 | | - | - |
| Tukey | Acq2 | test | | - | - |
| Tukey | Acq3 | test | | - | - |
| ANOVA | All sessions | D2ChR2 | | 17.03094193 | 4 |

| | | | | |
|---|---|---|---|---|
| Tukey | Pre-test | Acq1 | - | - |
| Tukey | Pre-test | Acq2 | - | - |
| Tukey | Pre-test | Acq3 | - | - |
| Tukey | Pre-test | test | - | - |
| Tukey | Acq1 | Acq2 | - | - |
| Tukey | Acq1 | Acq3 | - | - |
| Tukey | Acq1 | test | - | - |
| Tukey | Acq2 | Acq3 | - | - |
| Tukey | Acq2 | test | - | - |
| Tukey | Acq3 | test | - | - |
| ANOVA | All sessions | DArchT | 18.78309525 | 4 |
| Tukey | Pre-test | Acq1 | - | - |
| Tukey | Pre-test | Acq2 | - | - |
| Tukey | Pre-test | Acq3 | - | - |
| Tukey | Pre-test | test | - | - |
| Tukey | Acq1 | Acq2 | - | - |
| Tukey | Acq1 | Acq3 | - | - |
| Tukey | Acq1 | test | - | - |
| Tukey | Acq2 | Acq3 | - | - |
| Tukey | Acq2 | test | - | - |
| Tukey | Acq3 | test | - | - |

| | p-value | |
|---|---|---|
| - | 0.419823559 | |
| - | 0.419823559 | |
| - | 0.010680949 | # |

| df2 | p-value |
|---|---|
| 15 | 0.245780283 |

| df2 | p_value | |
|---|---|---|
| 16 | 0.00041515 | |
| | 0.002227006 | * |
| | 0.673037756 | |
| | 0.00079773 | |
| 16 | 0.011034716 | |
| | 0.159999642 | |
| | 0.241145508 | |
| | 0.008634788 | |
| 16 | 9.06977E-05 | |
| | 0.006672468 | * |
| | 0.050329326 | |
| | 7.00701E-05 | |
| 16 | 0.003546559 | |
| | 0.614093658 | |
| | 0.019212047 | * |
| | 0.003193655 | |
| 16 | 0.00089452 | |
| | 0.000747187 | * |
| | 0.023277393 | * |
| | 0.431211229 | |
| 30 | 0.493003014 | |
| - | 0.999998633 | |
| - | 0.956550603 | |
| - | 0.962548672 | |
| - | 0.850455532 | |
| - | 0.944928614 | |
| - | 0.971501746 | |
| - | 0.827609747 | |
| - | 0.658967079 | |
| - | 0.997779284 | |
| - | 0.466603097 | |
| 30 | 2.17591E-07 | |

| | Group1 |
|---|---|
| ANOVA | All sessions |
| ANOVA | All sessions |
| ANOVA | All sessions |
| ANOVA | All groups |
| ANOVA | All groups |
| ANOVA | All groups |
| ANOVA | All groups |
| ANOVA | All groups |

| | Group1 |
|---|---|
| ANOVA | All sessions |
| ANOVA | All sessions |
| | **Group1** |
| T-test | Laser |
| T-test | Laser |
| T-test | Laser |
| T-test | Laser |
| T-test | Laser |

| | Group1 |
|---|---|
| ANOVA | All sessions |
| ANOVA | All sessions |
| | **Group1** |
| T-test | Laser |
| T-test | Laser |
| T-test | Laser |
| T-test | Laser |
| T-test | Laser |

| | Group1 |
|---|---|
| ANOVA | All sessions |
| Post-hoc | Pre-test |
| Post-hoc | Pre-test |
| Post-hoc | Pre-test |
| Post-hoc | Pre-test |
| Post-hoc | Acq1 |
| Post-hoc | Acq1 |
| Post-hoc | Acq1 |
| Post-hoc | Acq2 |

| | | |
|---|---|---|
| - | 1.08444E-05 | # |
| - | 4.61026E-07 | # |
| - | 1.94543E-05 | # |
| - | 3.13221E-06 | # |
| - | 0.774096788 | |
| - | 0.999535062 | |
| - | 0.991021415 | |
| - | 0.651156646 | |
| - | 0.953069318 | |
| - | 0.963067465 | |
| 20 | 1.50069E-06 | |
| - | 0.027901172 | # |
| - | 6.03975E-05 | # |
| - | 0.000868125 | # |
| - | 0.808897449 | |
| - | 0.079890057 | |
| - | 0.540059531 | |
| - | 0.002486757 | |
| - | 0.750414652 | |
| - | 6.1011E-06 | |
| - | 7.6752E-05 | |

| | |
|---|---|
| Post-hoc | Acq2 |
| Post-hoc | Acq3 |
| ANOVA | All sessions |
| Post-hoc | Pre-test |
| Post-hoc | Pre-test |
| Post-hoc | Pre-test |
| Post-hoc | Pre-test |
| Post-hoc | Acq1 |
| Post-hoc | Acq1 |
| Post-hoc | Acq1 |
| Post-hoc | Acq2 |
| Post-hoc | Acq2 |
| Post-hoc | Acq3 |
| | **Group1** |
| T-test | Laser |
| T-test | Laser |
| T-test | Laser |
| T-test | Laser |
| T-test | Laser |

Fig 6e

| Group2 | Condition | F-value | df1 | df2 | p-value |
|---|---|---|---|---|---|
| D2tdTomato | HFD | 0.158973161 | 4 | 20 | 0.956587099 |
| D2ChR2 | HFD | 0.628110036 | 4 | 30 | 0.646185127 |
| D2ArchT | HFD | 1.239191514 | 4 | 25 | 0.319861017 |
| - | Pre-test | 3.460494511 | 2 | 15 | 0.058107656 |
| - | Acq1 | 0.124179258 | 2 | 15 | 0.88412004 |
| - | Acq2 | 0.172939402 | 2 | 15 | 0.842842086 |
| - | Acq3 | 0.334594706 | 2 | 15 | 0.720834335 |
| - | test | 1.469614352 | 2 | 15 | 0.261310098 |

Fig 6f

| Group2 | Condition | F-value | df1 | df2 | p-value |
|---|---|---|---|---|---|
| D2tdTomato | Laser | 0.529719247 | 4 | 20 | 0.715271197 |
| D2tdTomato | No Laser | 0.491580096 | 4 | 20 | 0.741943705 |
| **Group2** | **Condition** | **T-value** | **df** | | **p-value** |
| No Laser | Pre-test | 3.31658794 | 4 | - | 0.029472331 |
| No Laser | Acq1 | 0.99350249 | 4 | - | 0.37669958 |
| No Laser | Acq2 | -0.71197381 | 4 | - | 0.515804502 |
| No Laser | Acq3 | -0.14645233 | 4 | - | 0.89064881 |
| No Laser | test | 0.12724359 | 4 | - | 0.90488785 |

Fig 6 g

| Group2 | Condition | F-value | df1 | df2 | p-value |
|---|---|---|---|---|---|
| D2ChR2 | Laser | 0.952730061 | 4 | 30 | 0.447535722 |
| D2ChR2 | No Laser | 0.172208894 | 4 | 30 | 0.950916553 |
| **Group2** | **Condition** | **T-value** | **df** | | **p-value** |
| No Laser | Pre-test | -0.21606075 | 6 | - | 0.83609857 |
| No Laser | Acq1 | -1.26474046 | 6 | - | 0.252867213 |
| No Laser | Acq2 | -0.47941714 | 6 | - | 0.648615811 |
| No Laser | Acq3 | -0.50934505 | 6 | - | 0.628695361 |
| No Laser | test | 0.853988665 | 6 | - | 0.425887057 |

Fig 6 h

| Group2 | Condition | F-value | df1 | df2 | p-value |
|---|---|---|---|---|---|
| D2ArchT | Laser | 3.146853052 | 4 | 25 | 0.031699775 |
| Acq1 | Laser | - | - | - | 0.999759452 |
| Acq2 | Laser | - | - | - | 0.074168736 |
| Acq3 | Laser | - | - | - | 0.145226162 |
| test | Laser | - | - | - | 0.610036935 |
| Acq2 | Laser | - | - | - | 0.106352968 |
| Acq3 | Laser | - | - | - | 0.200411266 |
| test | Laser | - | - | - | 0.717016173 |
| Acq3 | Laser | - | - | - | 0.996940206 |

| | | | | | | |
|---|---|---|---|---|---|---|
| test | Laser | - | - | - | 0.685841945 | |
| test | Laser | - | - | - | 0.864505871 | |
| D2ArchT | No Laser | 4.148794583 | 4 | 25 | 0.010314067 | |
| Acq1 | No Laser | - | - | - | 0.96438896 | |
| Acq2 | No Laser | - | - | - | 0.029893118 | # |
| Acq3 | No Laser | - | - | - | 0.029791941 | # |
| test | No Laser | - | - | - | 0.403190996 | |
| Acq2 | No Laser | - | - | - | 0.119056491 | |
| Acq3 | No Laser | - | - | - | 0.118710727 | |
| test | No Laser | - | - | - | 0.787745364 | |
| Acq3 | No Laser | - | - | - | 0.99999999 | |
| test | No Laser | - | - | - | 0.643910937 | |
| test | No Laser | - | - | - | 0.642998372 | |
| **Group2** | **Condition** | **T-value** | **df** | | **p-value** | |
| No Laser | Pre-test | 1.653493142 | 5 | - | 0.159137566 | |
| No Laser | Acq1 | -0.74721104 | 5 | - | 0.488563803 | |
| No Laser | Acq2 | -4.40199017 | 5 | - | 0.007008818 | @ |
| No Laser | Acq3 | -4.3929974 | 5 | - | 0.007068018 | @ |
| No Laser | test | -1.91805023 | 5 | - | 0.113211354 | |
